# Bayesian analysis of longitudinal RB-TnSeq resolves the fitness seascape in fluctuating environments

**DOI:** 10.64898/2026.02.11.703069

**Authors:** Carl J. Stone, Megan G. Behringer

**Affiliations:** Department of Biological Sciences, Vanderbilt University, Nashville, TN 37232; Evolutionary Studies Initiative, Vanderbilt University, Nashville, TN 37232; Vanderbilt Institute for Infection, Immunology and Inflammation, Nashville, TN 37232; Department of Pathology, Microbiology and Immunology, Vanderbilt University Medical Center, Nashville, TN 37232

**Keywords:** starvation, quantitative genetics, trade-offs, *Escherichia coli*

## Abstract

Temporally structured environments are ubiquitous in nature, but time-dependent fitness effects are difficult to measure and thus understudied. To resolve temporal fitness structure at genome scale, we developed a Bayesian multilevel framework for longitudinal randomly barcoded transposon sequencing (RB-TnSeq) that stabilizes noisy mutant abundance trajectories into time-resolved selection-rate estimates with interpretable uncertainty. Applying this approach to a feast–famine starvation regime in *Escherichia coli*, we uncovered distinct fitness trajectories underpinned by shared molecular strategies across growth-curve phase, consistent with shifting constraints and antagonistic pleiotropy. Fitness during initial growth strongly constrained cumulative success, such that later advantages under stress could not rescue mutants that were initially deleterious. We then compressed these dynamics with a one-dimensional Fisher’s geometric “seascape” model that orders mutants along a latent axis aligning with generalist–specialist and growth–survival trade-offs, providing a compact quantitative description of genome-wide constraints in a fluctuating environment. Finally, our longitudinal estimates and inferred seascape are predictive of both the identity and timing of mutational targets in an extended evolution experiment under similar repeated feast–famine conditions, linking short-term competitive fitness effects, with their related trade-offs and constraints, to long-term adaptive outcomes.

## INTRODUCTION

Understanding how genetic variation shapes organismal fitness in fluctuating environments is a central challenge in evolutionary biology (Abdul-Rahman et al. 2021; Yamamichi et al. 2023). In natural environments, adaptation rarely occurs under constant conditions. These dynamic conditions are exemplified by microbial populations in the gut, soil, aquatic ecosystems, and host-associated niches which encounter repeated shifts in nutrient availability, stressors, and ecological interactions (Dethlefsen and Relman 2011; Thaiss et al. 2015; Woodhouse et al. 2016; Kram et al. 2017; Smits et al. 2017; Meisner et al. 2021; Behringer et al. 2022; Greenwald and Wolfgang 2022). Under such dynamics, variants that are beneficial in one condition may be neutral or deleterious in another, and their transient effects can determine the fate of lineages and the predictability of evolutionary trajectories (Nguyen et al. 2021; Behringer et al. 2024). Resolving these fine-scale, time-dependent contributions of individual mutations is therefore critical for linking genotypes to evolutionary outcomes.

High-throughput assays such as pooled CRISPR perturbation screens, barcoded lineage tracking, and transposon-insertion sequencing (TnSeq) have transformed the study of fitness landscapes by enabling genome-wide measurement of fitness effects for thousands of mutations in parallel (van Opijnen et al. 2009; van Opijnen and Camilli 2013; Shalem et al. 2014; Levy et al. 2015). These approaches have yielded genome-wide maps of genotype–phenotype relationships, quantified the role of clonal interference and epistasis, and illuminated the genetic basis of traits such as antibiotic resistance and host colonization (Goodman et al. 2009; van Opijnen et al. 2009; Gallagher et al. 2011; Levy et al. 2015; Venkataram et al. 2016; Li et al. 2023; Abreu et al. 2024; Couce et al. 2024). Yet, despite their power, these methods are largely restricted to static or endpoint measurements. The intrinsic noise in sequencing-based assays makes it especially difficult to resolve small or transient fitness effects with statistical confidence, leading to a methodological trade-off between genome-wide scale and temporal resolution (Chao et al. 2016). As a result, many fitness effects that may be central to adaptation in realistic, dynamic environments remain undetected.

Bayesian statistical methods provide a powerful framework to overcome these limitations. Bayesian inference can incorporate prior knowledge of neutral expectations and sequencing count distributions, producing posterior probability distributions that propagate uncertainty across time points. Historically, applying Bayesian inference at genome scale has been limited by computational difficulties due to the high-dimensional parameter space of high-throughput fitness assays; however, recent advances in statistical computing have lowered these barriers, making genome-wide Bayesian inference increasingly practical for noisy pooled assays (Zhang et al. 2022). As such, Bayesian approaches have been applied in similar high-throughput and longitudinal contexts, including assessing the fitness of novel variants in high-throughput lineage tracking assays and longitudinal analyses of metagenomic data (Sweeny et al. 2023; Razo-Mejia et al. 2024; Kim et al. 2025). However, Bayesian longitudinal inference has not yet been applied to Tn-Seq to resolve genome-scale, time-dependent fitness effects within fluctuating environments.

Bacterial starvation provides an ideal model for studying fitness in a fluctuating environment. In *Escherichia coli*, the transitions from exponential growth through stationary phase, death phase, and long-term stationary phase (LTSP) encompass reproducible shifts in nutrient availability, pH, resource scavenging, and stress responses (Zambrano et al. 1993; Finkel and Kolter 1999; Zinser and Kolter 2000; Behringer et al. 2018; Behringer et al. 2022). These dynamic selection pressures across growth phases, coupled with the inherent noise in barcode-based measurements, have limited the ability to use pooled mutant libraries to resolve transient or subtle fitness effects. As such, investigation of the genes that contribute to fitness across the microbial growth curve has largely focused on large-effect global regulatory pathways—such as sigma factor regulons, the stringent response, and catabolite repression—but the combined effect of transient contributions of individual genes across the growth curve remain underexplored (Zambrano et al. 1993; Zinser and Kolter 2000; Gosset et al. 2004; Nair and Finkel 2004; Kram et al. 2020; Irving et al. 2021).

Here, we develop a Bayesian framework for longitudinal inference of mutant fitness from randomly barcoded TnSeq (RB-TnSeq) data. Applying this approach to an *E. coli* library sampled over a 10-day starvation regime, we obtain accurate, interpretable confidence intervals for thousands of selection-rate estimates. This allows us to detect subtle, condition-specific fitness effects that are obscured by standard endpoint analyses. We identify distinct temporal trajectories associated with stress-response regulation, biofilm modulation, and resource scavenging. Then, we compress these time-resolved fitness dynamics into a one-dimensional Fisher’s geometric “seascape” model, providing a compact description of genome-wide trade-offs across growth, death, and LTSP. Finally, we show that our results parallel patterns observed in long-term evolution experiments under similar selective regimes (Kram et al. 2017; Behringer et al. 2018; Katz et al. 2021; Behringer et al. 2024). Together, these findings demonstrate how rigorous statistical modeling can unlock fine-scale maps of gene–fitness relationships in fluctuating environments, advancing both functional genomics and evolutionary biology.

## RESULTS

### Bayesian modeling improves the accuracy and interpretability of longitudinal RB-TnSeq

To obtain time-resolved fitness estimates from longitudinal RB-TnSeq while retaining interpretable uncertainty, we developed a Bayesian framework that infers gene-by-interval selection rates from barcode trajectories and returns posterior distributions for each estimate. We validated this approach by benchmarking it against a widely-used *t*-test—based analysis method (BarSeq) on simulated longitudinal data (**Fig. S1**) (Wetmore et al. 2015). Simulations incorporated varying parameters for likely sources of experimental noise: count variability between different transposon insertions in the same gene, random measurement error (a single parameter for multiple possible stochastic processes like amplification bias, differences between replicates, etc.), and directional batch effects. We chose noise parameter ranges similar to the magnitude of within-gene, between-replicate, and between-batch variability we have observed in RB-TnSeq measurements (**Fig. S2**). Across all simulations, our model achieved accuracy comparable to BarSeq in estimating true selection rates (**Fig. S1A**; our model: median RMSE 0.41, 95% CI [0.39, 0.44]; BarSeq: median RMSE 0.40, 95% CI [0.38, 0.43]).

Along with similar accuracy to BarSeq, our model consistently exhibited higher coverage probability, defined as the fraction of credible intervals containing the true value (**Fig. S1B**; our model: median coverage 53%, 95% CI [49%, 57%]; BarSeq: median coverage 39%, 95% CI [33%, 42%]). This advantage was most pronounced under high within-gene variability and high measurement error, where coverage increased to 58% (95% CI [53%, 65%]) compared to 23% (95% CI [18%, 29%]) for BarSeq (**Fig. S1D**). Large batch effects reduced coverage probability in both methods. Overall, these simulations indicate that our Bayesian model preserves accuracy while substantially improving uncertainty quantification in longitudinal RB-TnSeq experiments, especially under the noisy conditions characteristic of sequencing-based assays like Tn-Seq.

### Longitudinal fitness analysis resolves shifts in adaptive pressure during the entry into long-term stationary phase

After validating our model, we applied it to an RB-TnSeq library in *E. coli* to quantify genome-wide, gene-specific fitness effects across the extended microbial growth curve. For each interval between sampling points, the model estimates a posterior distribution for the selection rate of each gene relative to an empirically neutral gene set, defined as genes whose mutants have consistently near-zero selection rates across a large compendium of growth and stress conditions (**Table S1**) (Price et al. 2018). These interval-specific selection rate estimates (denoted for each interval 𝑡 → 𝑡 + 1 as 𝑠_𝑡→𝑡+1_) are inferred simultaneously across all three intervals which can then be combined as a time-weighted average to calculate the net selection rate over the full experiment (𝑠_𝑛𝑒𝑡_; **Fig. 1A**). Because inference produces posterior distributions rather than point estimates, uncertainty can easily be propagated into downstream analyses.

**Figure 1.**
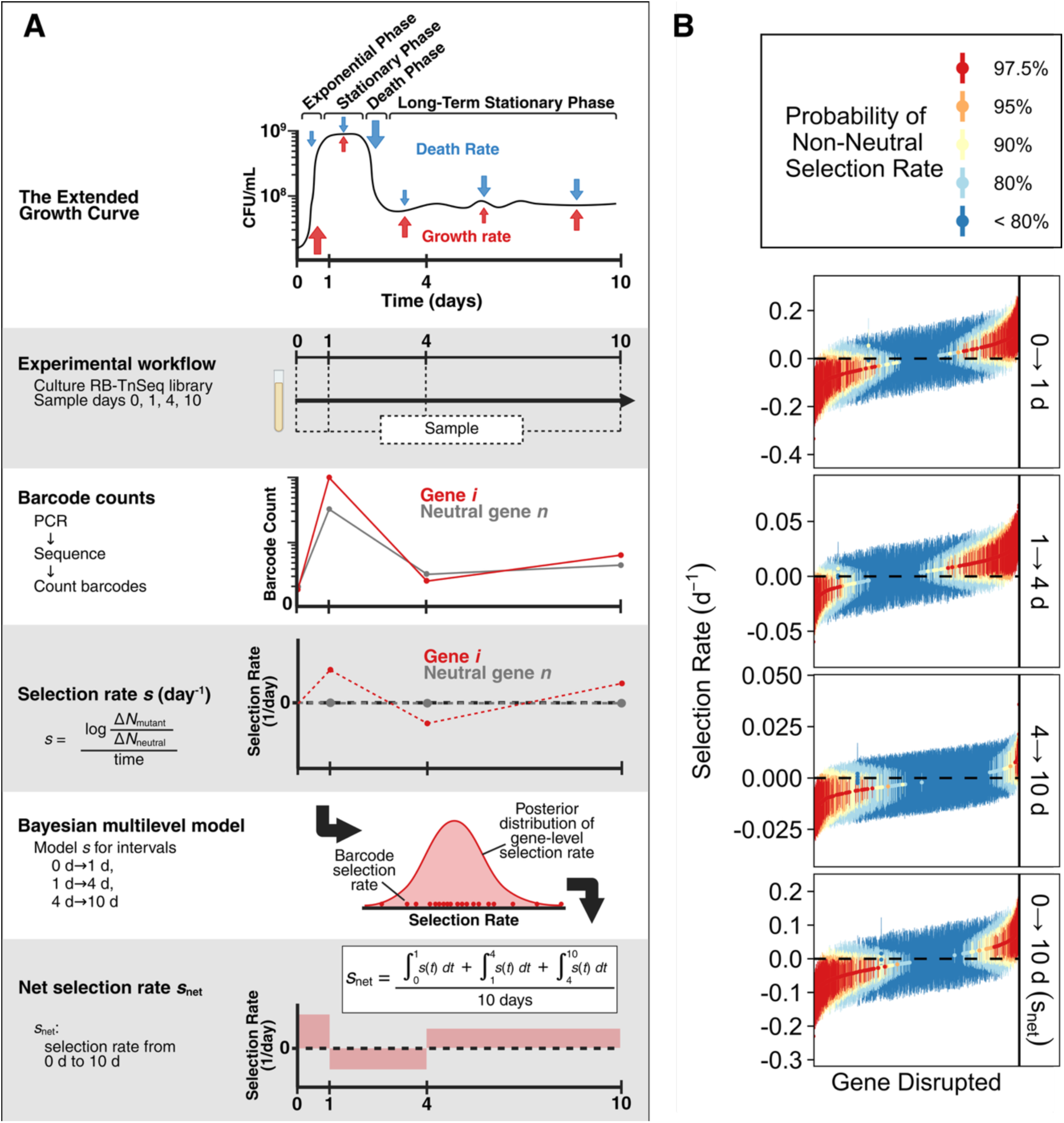
Inference of transposon mutant selection rates over extended culture. **(A)** Schematic of the experimental workflow and longitudinal Bayesian model used to infer gene-level selection rates across sequential time intervals. **(B)** Posterior summaries of gene-level selection rates for each interval (0→1 d, 1→4 d, 4→10 d) and for the average effect across the full experiment (𝑠_𝑛𝑒𝑡_). Each point and vertical line shows the posterior median and 95% credible interval for one gene. Colors show the posterior probability that the selection rate is positive or negative (i.e. Pr(s > 0) or Pr(s < 0)). Genes are ranked within each panel by median selection rate, and the gene order is independent across panels.

We applied this framework to the *E. coli* transposon library grown for 10 days in LB medium. The barcoded transposon-mutant library was sampled immediately after inoculation (day 0) and again at days 1, 4, and 10, providing a time series that captured mutant abundance trajectories across the major physiological transitions of the extended growth curve: rapid growth, stationary phase, death phase, and lastly long-term stationary phase (Zambrano et al. 1993; Finkel 2006). Selection rates were estimated for each time interval (𝑠_0→1_, 𝑠_1→4_, and 𝑠_4→10_) relative to the neutral gene set. We identified 822 genes that were annotated by Wetmore *et al*. as present in the RB-TnSeq library but were not detected in our samples (**Table S2**) (2015). These genes are essential or highly deleterious when disrupted and were likely purified from the library before we acquired it or lost during preparatory cultures (Wetmore et al. 2015). Functional enrichment analysis revealed that these genes are largely associated with ribosome biogenesis, aminoacyl-tRNA synthesis, lipid metabolism, DNA replication and repair, and one-carbon metabolism (**Fig. S3**).

Out of the 3668 assessed genes present in our library, we then identified 1067 genes whose disruption conferred confidently positive or negative fitness effects in at least one time interval (**Table 1**, **Fig. 1B**). Mutants were classified as non-neutral if their posterior probability distributions indicate at least a 90% probability that their selection rate is above or below zero. Fitness distributions varied considerably across time intervals (**Table 1**, **Fig. S4, Fig. S5**). Mutations were slightly biased toward deleterious effects during the interval 0→1 d, then were showed more beneficial effects from 1→4 d and more deleterious effects from 4→10 d. These shifts illustrate that selective pressures change dramatically as environmental conditions fluctuate over the course of starvation. We then averaged interval-specific selection rates to calculate 𝑠_𝑛𝑒𝑡_, the cumulative selection rate of each mutant over the 10-day experiment. Most gene disruptions (733 genes) were cumulatively deleterious, although a subset (361 genes) were cumulatively beneficial across the 10 days (**Table 1**, **Dataset S1, Fig. 1B**, bottom panel; Fisher-Freeman-Halton test: 𝑝 = 5.0 × 10^−6^, 𝑠_𝑛𝑒𝑡_ 𝜒^2^ goodness-of-fit with BH correction: 𝑝 = 5.4 × 10^−22^). Using a stricter probability cutoff of greater than 97.5% reduced the number of confidently non-zero mutant fitness effects but did not qualitatively change any of the temporal or cumulative fitness trends (**Table S3**).

**Table 1.**
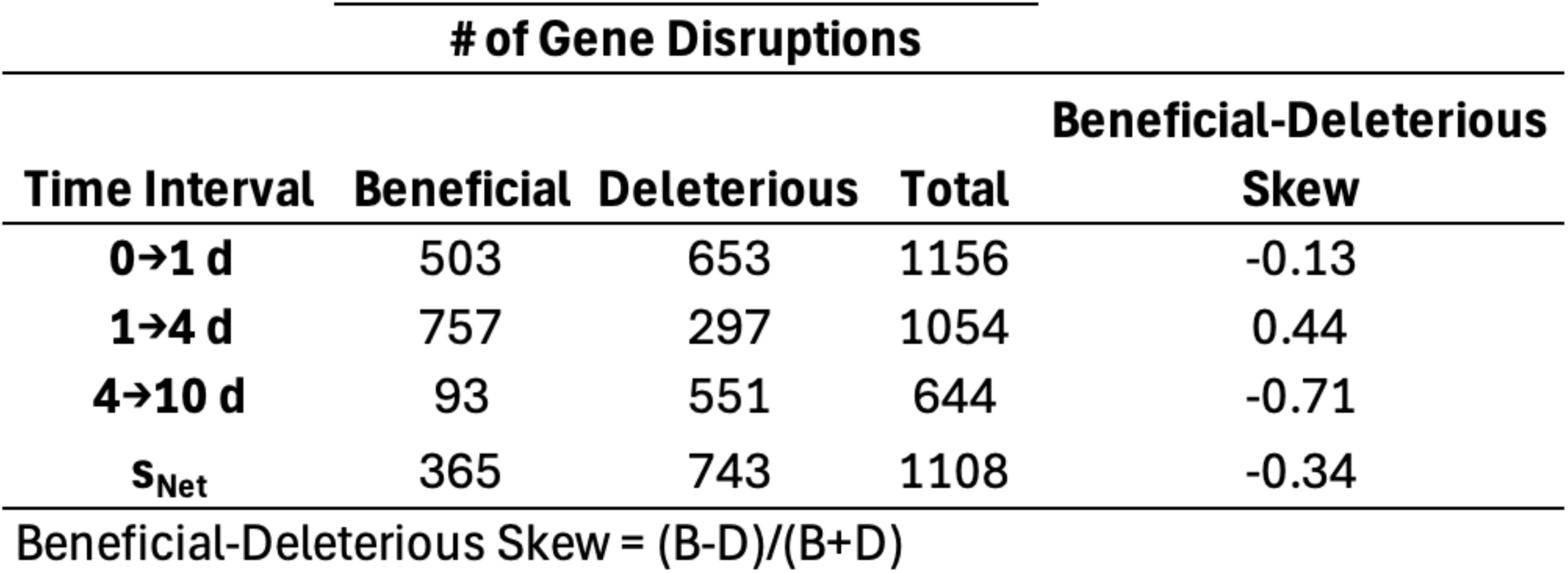
Number of confidently non-zero transposon mutants by time interval. Genes were classified as beneficial or deleterious when the posterior probability that the gene’s selection rate was above or below zero exceeded 90% (i.e., Pr(s > 0) > 0.9 or Pr(s < 0) > 0.9).

### Gene clusters reveal distinct temporal fitness strategies

Next, we grouped non-neutral genes by their selection rate trajectories to identify temporal fitness trade-offs. Principal component analysis (PCA) of selection rates across the three intervals revealed that the first principal component captured 80% of the variance and aligned with the combined eigenvector of 𝑠_0→1_ and 𝑠_4→10_ while being strongly anticorrelated with 𝑠_1→4_, which spans death phase, during which culture cell counts decrease by ∼10-100—fold (Pearson correlation of selection rates: r_0→1:1→4_ = −0.90, *p* < 2.2×10^−16^; r_1→4:4→10_ = −0.76, *p* < 2.2×10^−16^) (Finkel 2006). This anticorrelation between adjacent time intervals is also present in the raw selection rates that were inputs to our Bayesian model (r_0→1:1→4_ = −0.41, *p* < 2.2×10^−16^; r_1→4:4→10_ = −0.44, *p* < 2.2×10^−16^). The second principal component (20% of variance) primarily separated exponential-phase effects from LTSP effects, distinguishing mutants who were more beneficial in 0 → 1d from those that were beneficial in 4 → 10d. We then grouped mutants into distinct fitness trajectories using k-means clustering on each mutant’s three interval selection rates (k = 4, **Fig. S6**). Mutants within clusters represent distinct survival strategies across long-term culture (**Fig. 2A**). The resulting clusters had similar sizes ranging from 329 genes in cluster IV to 432 genes in cluster I (**Table S4**). Each cluster was organized by the sign of 𝑠_0→1_ and 𝑠_4→10_ to facilitate biological interpretation (**Fig. 2B-C**). Interestingly, only clusters I and IV, both of which included mutants with positive 𝑠_0→1_, contained genes with positive 𝑠_𝑛𝑒𝑡_. In contrast, although cluster II contained mutants with beneficial effects at day 10, these effects were not strong enough to overcome initial deleterious effects during the 0→1 d interval. This suggests that early fitness exerts a dominant influence on cumulative success over this culture length.

**Figure 2.**
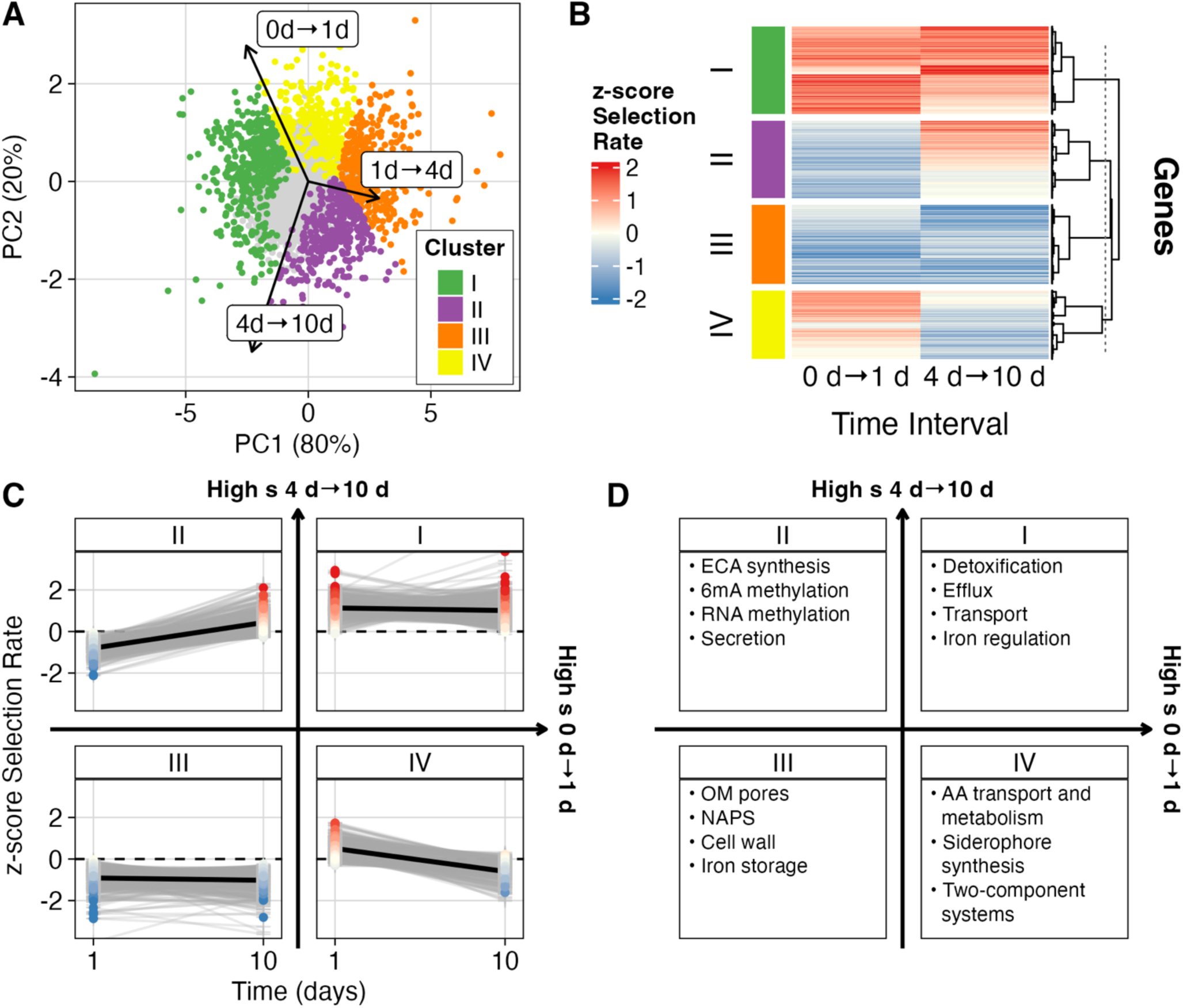
Tn mutants cluster into distinct temporal selective trajectories. **(A)** Principal component analysis of gene-level selection rate estimates across time intervals. Arrows indicate the loading of each interval on the two PCs. Each point represents one gene. Genes with non-zero selection rates in intervals 0→1 d or 4→10 d are colored, and genes with no confidently non-zero estimates are gray. Non-neutral genes were clustered by cosine distance, with cluster assignment shown by color. **(B)** Heatmap of z-scored selection rate estimates of early versus late selection (0→1 d and 4→10 d). All values outside [−2, 2] are clipped to those boundaries for visualization. Genes are clustered by selection rate similarity. **(C)** Cluster-specific temporal trajectories. Each gray line connects a single gene’s scaled selection rate estimates across intervals. Panel labels indicate the cluster identity. Black lines show a robust linear fit summarizing the cluster trend. Arrows between panels form a coordinate plane indicating the relative order of clusters by selection rate at two intervals: horizontal axis = low 0→1 d to high 0→1 d; vertical axis = low 4→10 d to high 4→10 d. **(D)** Functional categories overrepresented among genes in each cluster based on annotations from Gene Ontology, KEGG, and EcoCyc pathways (FDR q < 0.1). As in C, panel labels indicate the cluster identity, and arrows indicate the relative order of clusters by selection rate at early and late intervals.

To investigate the biological functions associated with each fitness trajectory, we performed functional enrichment analysis using gene ontology terms, KEGG pathways, and EcoCyc pathway annotations (**Fig. 2C, Table S5**) (Ashburner et al. 2000; Kanehisa and Goto 2000; Kanehisa 2019; Kanehisa et al. 2025; Karp et al. 2025; The Gene Ontology Consortium 2026). Cluster I, which showed consistent benefits early and late in culture, was enriched for genes encoding efflux systems, particularly those associated with detoxification (**Fig. 2D**). Cluster II, which gained fitness late in culture, contained mutants related to the biosynthesis of enterobacterial common antigen, secretion of surface structural components such as pili and flagella, and multiple methyltransferases, including *dam* and several involved in rRNA and tRNA modification. These genes are important for growth, metabolism, and membrane homeostasis, making them deleterious to mutate early but dispensable, and possibly energetically favorable to lose, at later times.

Cluster III genes were consistently deleterious when disrupted, suggesting roles in critical processes. This group included genes involved in central metabolism, outer membrane integrity, nucleoid organization, and iron storage. In contrast, cluster IV mutants were initially beneficial but had neutral or negative 𝑠_4→10_ and were enriched for genes involved in conditional stress responses, such as amino acid transport, two-component systems, and siderophore synthesis. The loss of these genes may confer short-term energetic advantages that become costly as the environment changes.

In addition to identifying functional enrichment across clusters, we examined the mutants with the largest fitness effects to identify major physiological strategies that shape survival and competition during prolonged culture. Efflux systems showed particularly strong and coherent patterns in 𝑠_𝑛𝑒𝑡_ (**Fig. S7A**). Most individual efflux pumps were slightly beneficial when inactivated, whereas disruption of *mdtABC* was mildly deleterious, and disruption of *tolC*, required for all major efflux systems, was strongly deleterious (Zgurskaya and Nikaido 2000; Tikhonova et al. 2011). These patterns indicate that while pumps are partially redundant, a degree of efflux capacity remains essential during prolonged culture. The strongest positive 𝑠_𝑛𝑒𝑡_observed in this experiment was disruption of the repressor *cytR* (**Fig. S7B**) (Shin et al. 2001; Browning and Busby 2004). Consistent with *cytR* disruption being strongly beneficial, disruptions of genes in the *cytR* regulon were on average deleterious, and the regulon had more deleterious genes than the null expectation from size-matched random gene sets, where the expected number is calculated as the sum of each gene’s probability of being deleterious (Pr[*cytR* regulon > null] = 0.93; **Fig. S8**). Together these data suggest that derepression of the *cytR* regulon may provide a net advantage during starvation.

Biofilm-associated mutants also exhibited strong and time-specific fitness effects. Type I fimbriae mutants showed concerted and contrasting selection effects: activator *fimB* was deleterious and inactivator *fimE* was beneficial across both early and late intervals, indicating strong selection on fimbrial switching dynamics (**Fig. S7C**) (Klemm 1986). Curli fiber genes exhibited transient fitness effects: structural and export gene mutants (*csgA*, *csgE*) were deleterious early, while folding and polymerization gene mutants (*csgC*, *csgF*) were beneficial in the same time interval, but all curli-associated mutants were neutral at the later interval (Chapman et al. 2002). Similarly, the phase-variable adhesin Ag43 (*flu*) mutant had strong early benefits but became neutral later (Henderson et al. 1997; Wallecha et al. 2002). Together, these strong pathway-level effects exemplify the major physiological strategies shaping growth and survival, including detoxification, regulatory adaptation, and modulation of surface structures, and illustrate how time-resolved fitness measurements can identify discrete mechanistic contributors to long-term competition.

### A geometric seascape model recapitulates the growth-survival trade-off

Despite these strong pathway-level effects, these examples do not explain a broader model for how selection acts on the population. Mutants in disparate pathways show coordinated gains and losses of fitness between intervals, suggesting that selection is acting on a shared physiological trade-off rather than on a specific molecular function. To summarize this structure, we fit a one-dimensional Fisher’s geometric “seascape” model that assigns each transposon mutant genotype (𝑔) a latent phenotypic value (𝑧_g_, a unitless coordinate) and estimates a separate single-peaked fitness curve for each time interval (𝑑) (Fisher 1930; Tenaillon 2014). We call 𝑧 a latent phenotypic axis because it represents a multidimensional phenotypic state that is theoretically measurable but practically must be inferred from fitness. In this model, each interval’s landscape is defined by a fitness optimum at a fixed latent phenotypic value 𝑧 and a steepness (or curvature) that quantifies how strongly fitness declines as mutants deviate from the optimum (**Fig. 3A**, full specification in **Supplemental Text 1**). We fit the seascape model jointly to all mutants and time intervals, incorporating uncertainty from the longitudinal model, estimating a single latent coordinate for each gene and resolving a distinct landscape for each phase of the extended growth curve.

**Figure 3.**
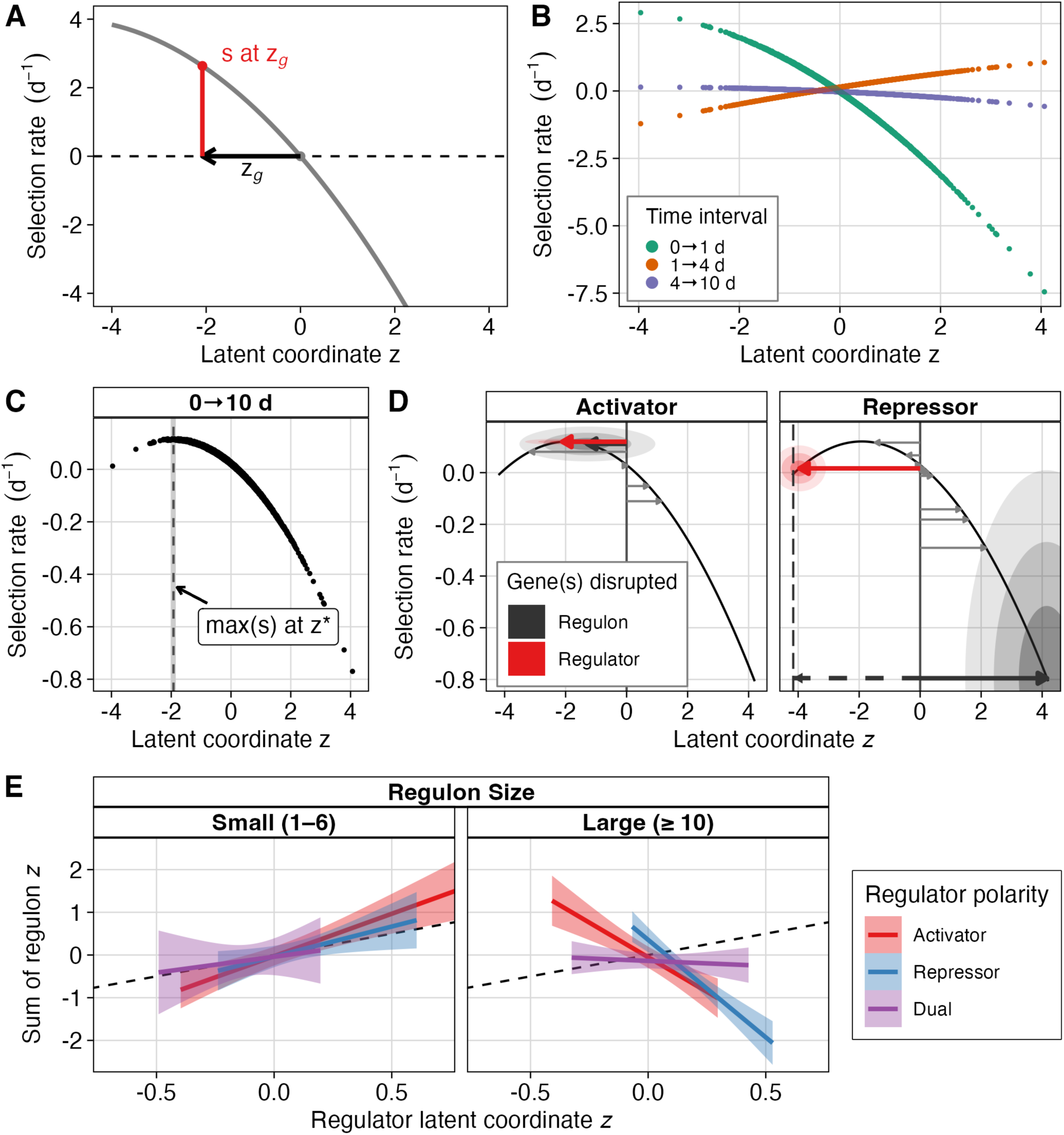
A geometric seascape model orders genes along a latent growth-survival axis. **(A)** Schematic of the one-dimensional Fisher’s geometric seascape. Each gene *g* is assigned a latent phenotypic coordinate 𝑧_g_, and the expected selection rate in each interval is modeled as a quadratic function of distance from the interval-specific optimum. **(B)** Interval-specific seascapes fitted to gene-level selection-rate posteriors from the longitudinal RB-TnSeq model. Points show the posterior median of gene-level selection rate estimate as predicted by the seascape model, with colors indicating the time interval. **(C)** Time-weighted integrated seascape across 0-10 d, computed as the time-weighted average of interval-specific seascapes. The integrated optimum 𝑧^∗^indicates the phenotypic coordinate that maximizes cumulative expected selection rate. **(D)** Phenotypic coordinates of activator *ydcI* mutant, repressor *cytR* mutant, and their regulon mutants. Red arrows show the 𝑧_g_ vector of regulator mutants of *ydcI* (left panel) and *cytR* (right panel), with concentric shaded ellipses showing credible intervals (center: 50%, middle: 80%, outside: 95%) for both 𝑧 and estimated selection rate. Gray arrows show the 𝑧_g_ vector for each mutant in the regulon. Black arrows show the signed sum of regulon mutants, with shaded ellipses showing credible intervals (50%, 80%, 95%). The dashed black arrow and vertical dashed line indicate the negative signed sum of the *cytR* regulon mutants. **(E)** The signed sum of regulon mutants correlates with regulator phenotypes in small regulons, but correlation decreases with regulon size.

The fitted seascape recapitulates the antagonistic selection pressures inferred from our interval-specific analyses (**Fig. 3B**). Selective pressure shifts in both magnitude and sign across our experimental intervals. The inferred fitness optima for early growth (0→1 d) and LTSP (4→10 d) lie on the far-negative end of the latent axis, whereas the death-phase (1→4 d) optimum is on the opposite, positive end. This suggests that there is a growth–survival trade-off between exponential phase and death phase that aligns LTSP more closely with growth than survival, consistent with repeated subpopulation turnover and growth on recycled resources (secreted metabolites, necromass, etc.) during LTSP (Finkel 2006). The interval-specific landscapes also vary in steepness, with selection being strongest early and decreasing monotonically over subsequent intervals. Combining the interval-specific functions yields the integrated 0→10 d landscape, which exhibits a single optimum 𝑧^∗^ (**Fig. 3C**). Mutants with 𝑧_g_ close to optimum 𝑧^∗^maximize cumulative growth and survival across the entire experiment. By contrast, mutants at extremely low or high 𝑧_g_ incur large costs during at least one stage of the extended growth curve. Altogether, this seascape allows us to order all transposon mutants in our library on the same latent axis, giving us a structured look into the interplay between genotypes, phenotypes, and selection in this shifting environment.

Using the latent axis from the seascape model, we investigated whether the 𝑧_g_ positions of regulators could be predicted from the 𝑧_g_ of mutants in their regulons. If a regulators latent coordinate primarily reflects the combined effects of its targets, then the regulator should lie near the signed sum of its targets on the 𝑧 axis; in other words, the 𝑧_g_ of activators should equal the sum of their targets’ 𝑧_g_, and the 𝑧_g_ of repressors should equal the negative sum of their targets. This expectation holds in specific cases: for activator *ydcI*, 𝑧_𝑦𝑑𝑐𝐼_is −2.11 [−3.32, −0.89] (median [95% CI]) and the sum of its regulon 𝑧_g_ is −1.29 [−3.53, 0.92]; for repressor *cytR*, 𝑧_𝑐𝑦𝑡𝑅_is −3.96 [−4.83, −3.10] while the sum of its regulon is 4.15 [1.45, 6.80] (**Fig. 3D**). This correlation is strongest for regulators with small regulons, consistent with a geometric model assumption that latent axis positions are additive; however, as regulon size increases, the relationship weakens and ultimately reverses, suggesting a heuristic boundary beyond which large, pleiotropic regulons reorganize physiology in ways that are not well-approximated by a one-dimensional model (**Fig. 3E**). These results highlight how network complexity can produce unexpected mappings between genetic perturbations and latent phenotypes.

To interpret the physiological meaning of the latent phenotypic axis, we compared seascape 𝑧_g_ coordinates to previously published gene-level fitness measurements across 131 conditions using the same RB-TnSeq library (**Table S6**) (Price et al. 2018). We first calculated the distance between fitness profiles for each condition then clustered conditions by similarity (**Fig. 4A**). Condition clusters split broadly into two groups: clusters 1, 2, and 3 comprised only nutritional assays where populations were grown in minimal media with either ammonium and a varying carbon source (hereafter, “single-carbon” experiments) or glucose and a varying nitrogen source (“single-nitrogen” experiments); clusters 4, 5, and 6 included diverse chemical and physical stresses (including antibacterials, metals, solvents, general biocides, and inorganic toxins), along with a small number of single-carbon conditions that grouped with stresses, particularly L-lactate (**Fig. 4B**, **Table S7**). We then correlated gene-level selection rates with seascape coordinates for each condition using a generalized additive model (**Fig. 4C, Fig. S9**). Genes at strongly negative 𝑧_g_ were near-neutral across all conditions. However, selection rate diverged across condition clusters as 𝑧_g_increased: high-𝑧_g_genes showed strongly positive selection rate in the most stressful conditions (cluster 6), but neutral to mildly deleterious selection rates in moderate stress (clusters 4 and 5) and strongly deleterious effects in growth on varying carbon or nitrogen sources (clusters 1, 2, and 3) (**Fig. 4D**). Thus, ordering genes along 𝑧 reveals a continuum from environmentally robust, weak-effect mutants to mutants with a strong trade-off favoring stress resistance at the expense of metabolic flexibility.

**Figure 4.**
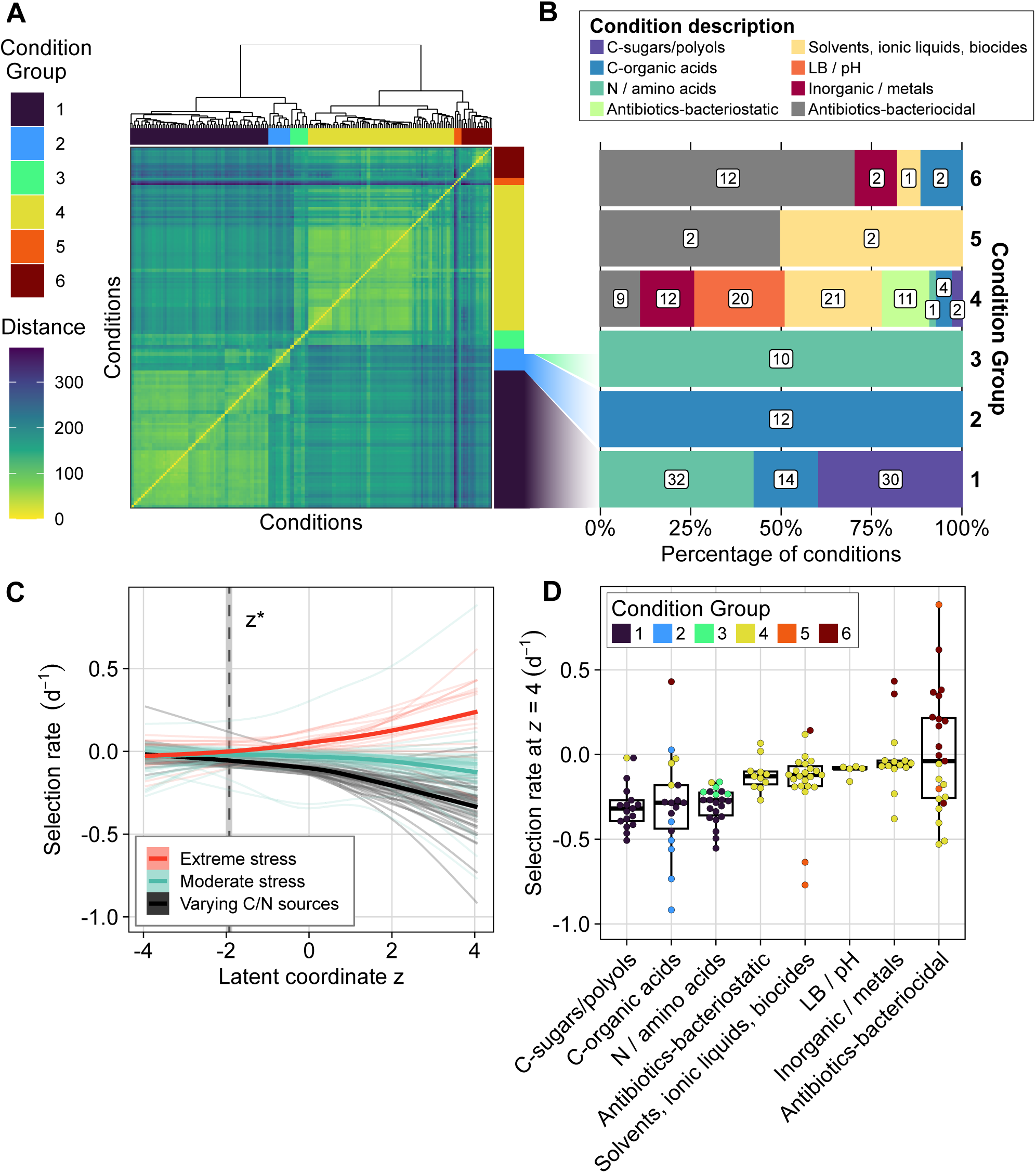
Mapping a published fitness compendium onto the seascape links the latent axis to growth-stress trade-offs across environments. **(A)** Hierarchical clustering of conditions from a published RB-TnSeq compendium using distance between genome-wide fitness profiles shows distinct groupings of experimental conditions. Conditions are shown clustered against each other on both axes, with hierarchical clustering indicated by the dendrogram. **(B)** Composition of each condition cluster by coarse condition class shows that clusters separate nutrient limitation from diverse stresses. **(C)** Comparison of fitness across environments to the seascape phenotypic axis shows extreme trade-offs between stress resistance and nutritional competence at high phenotypic values. Each line indicates the smoothed average selection rate in one experimental condition across the latent phenotypic axis, estimated using a generalized additive model. Thick lines (one for each of the three groups) indicate the average selection rate of all experimental conditions in that group. Experimental conditions are grouped as varying C/N sources (clusters 1, 2, and 3), moderate stress (clusters 4 and 5), and extreme stress (cluster 6). **(D)** Selection rate of conditions at the positive extreme of latent axis (*z* = 4), grouped by condition type and cluster. Each point represents one experiment. Selection rate values are predicted from the generalized additive models in panel C.

### Transposon mutant fitness correlates with positive selection in extended evolution experiments

To assess the ecological relevance of predicted fitness values from our Bayesian model, we compared our data to a previous experimental evolution study in which *E. coli* populations underwent repeated feast/famine cycles every 10 days for a total of 300 days (Behringer et al. 2022). Mutations of low frequency were discarded as spurious, and duplicate alleles resulting from mapping errors were manually removed. Alleles were then classified by mutation type and by the fitness of each Tn mutant in our library.

For each mutation type, a permutation test was conducted to establish a null expectation for the Tn fitness effect (**Fig. 5A**). Genes that showed high fitness upon disruption in our Tn screen exhibited elevated mutation acquisition rates across all mutation classes, including nonsynonymous, synonymous, nonsense, indel mutations, and even large insertions and deletions, though large indels had too few occurrences for reliable null distribution estimates. Genes identified as deleterious in our Tn fitness assay exhibited nonsynonymous and nonsense mutation rates consistent with null expectations. However, both genes with beneficial and deleterious Tn mutant effects showed notably higher rates of indels than expected, with beneficial genes exhibiting the highest rates. Interestingly, synonymous mutations occurred at increased rates in beneficial genes and decreased rates in deleterious genes compared to expectations, although the magnitude of the reduction in deleterious genes was modest yet statistically significant (**Fig. 5A**).

**Figure 5.**
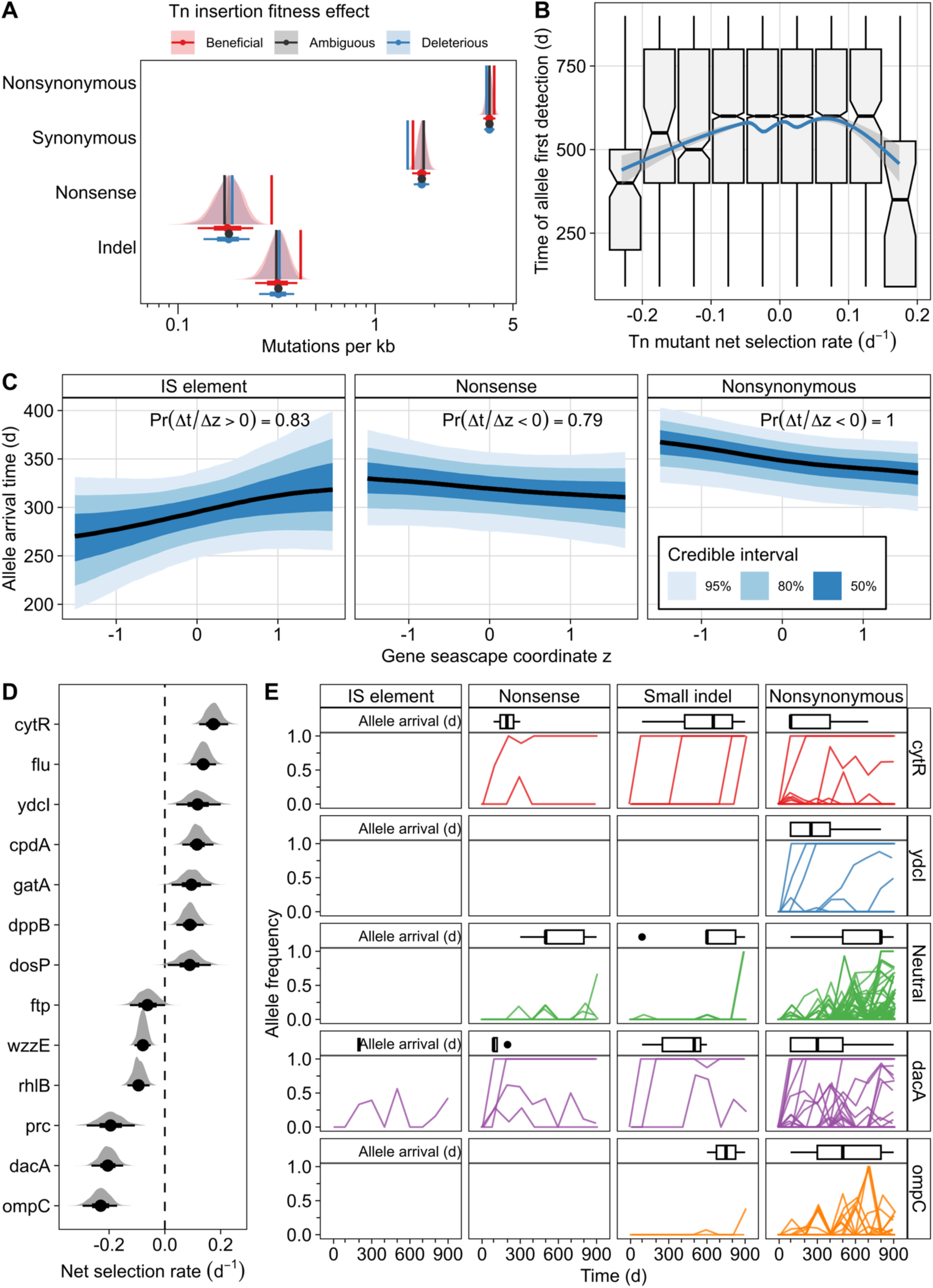
Transposon mutant fitness associates with mutational targets and timing in feast/famine experimental evolution. **(A)** Genes with strong transposon-mutant fitness effects acquire varying numbers of mutations during a repeated 10-day feast/famine evolution experiment. Distributions and interval bars summarize the null expectation for the number of genes mutated in each fitness bin (median, 80% and 95% intervals as drawn), and vertical lines show the observed number of mutations. **(B)** Mutations in genes with large-magnitude transposon fitness effects arise earlier than genes with small effects over repeated feast/famine evolution. Arrival time is summarized across replicate populations. The relationship between arrival time and selection rate was modeled as a GAM with a penalized cubic spline (blue line with 95% confidence interval). **(C)** Seascape axis coordinates correlate with arrival time of IS element insertions, nonsense mutations, and nonsynonymous mutations in populations adapted to feast/famine. Curves and ribbons are posterior summaries from a cumulative-logit ordinal mixed model predicting allele arrival time from a penalized spline of *z*. Ribbons show the indicated credible intervals. For each mutation class, predictions were compared at low versus high *z* (2.5^th^ vs 97.5^th^ percentile), and the posterior probability of a directional difference between *z* values is shown. **(D)** Transposon mutant fitness values for genes repeatedly mutated across multiple long-term starvation or feast/famine evolution studies show recurring targets can have strong beneficial or deleterious transposon mutant effects. **(E)** Examples of locus-specific mutation dynamics across evolution show distinct trajectories by mutation type. Trajectories show allele frequency time series for representative genes and mutation classes. Inset summaries show the distribution of first detection times for each unique allele. The “neutral” trajectories are mutations acquired in all empirically neutral reference set genes combined. Allele frequencies are colored by gene (or neutral genes) or visualization.

We next examined how mutation timing related to Tn mutant fitness. We found that mutations in genes with extreme Tn fitness values (𝑠_𝑛𝑒𝑡_ < −0.15 or > 0.15) arose, on average, approximately 100 days earlier than mutations in genes with intermediate fitness values (**Fig. 5B**). This suggests that absolute Tn fitness effect, rather than directionality, correlates with mutation timing across all types of mutations. However, the seascape axis showed clear trends between gene 𝑧_g_ coordinates and the arrival time of some types of mutations (**Fig. 5C**). IS-element insertions arose substantially earlier in genes with negative 𝑧_g_ than in genes with positive 𝑧_g_ . In contrast, nonsense and nonsynonymous mutations showed the opposite pattern, with earlier arrival in genes at higher 𝑧_g_. All other mutation classes showed no strong relationship between 𝑧_g_ and arrival time (**Fig. S10**). Together, these patterns indicate that the seascape captures temporal structure relevant to repeated adaptation in a similar growth regime, with the strongest predictive signal for the same type of alleles used to infer the seascape.

Further analysis concentrated on genes consistently targeted by mutations across multiple LTSP experimental evolution studies, including those with shorter feast/famine cycles (∼7-10 days) and prolonged stationary phases (hundreds of days to years) (**Fig. 5D**) (Kram et al. 2017; Behringer et al. 2018; Katz et al. 2021; Ratib et al. 2021; Behringer et al. 2022; Behringer et al. 2024). These repeatedly mutated genes exhibited a broad range of Tn mutant fitness effects, including genes with the highest and lowest 𝑠_𝑛𝑒𝑡_ in our library. Notably, specific temporal mutation dynamics emerged: *cytR* mutations, both nonsense and nonsynonymous, tended to sweep through populations within the first 90 days, followed by small indels that fixed around day 600 (**Fig. 5E, Fig. S11**). Contrasting mutation dynamics were observed in *dacA* and *ompC*, both highly deleterious in our Tn assay. *dacA* rapidly accumulated IS elements, nonsense, nonsynonymous mutations, and small indels early in evolution. In contrast, *ompC* showed no fixation events but accumulated numerous nonsynonymous mutations and indels that fluctuated without fixation over time (**Fig. 5E**).

Collectively, these findings underscore that Tn mutant fitness effects can substantially differ from those of other mutation types, even presumed inactivating mutations. Nonetheless, Tn mutant fitness may serve as a meaningful indicator of adaptive targets, highlighting genes likely to accumulate mutations in highly selective environments. Moreover, mutation timing followed mutation-type-specific trends along the latent seascape axis, showing that this approach can predict adaptation in mutations similar to the seascape dataset but also captures associations in unrelated mutation types that can be investigated further. Although inactivating mutations correlated moderately with Tn fitness, the timing of mutation acquisition displayed a stronger association with the absolute magnitude of Tn fitness effects than the direction of the effect.

## DISCUSSION

Here, we present a robust framework for analyzing barcoded high-throughput fitness assays in fluctuating environments. RB-TnSeq has been transformative for mapping gene-fitness relationships across diverse stresses, but technical limitations have typically limited analyses to endpoint readouts. By pairing longitudinal sampling with Bayesian inference, we have stabilized noisy fitness trajectories, obtained interpretable posterior credible intervals for time-dependent selection rates, and propagated uncertainty into downstream analyses. Applied to the extended *E. coli* growth curve, this approach resolves transient and cumulative fitness effects genome-wide, revealing structured trade-offs across the growth–death–LTSP transition (Finkel and Kolter 1999; Kram et al. 2020). Our results align late stationary-phase selective pressures with earlier growth phase and highlight loci that repeatedly shape evolutionary outcomes under feast/famine starvation regimes (Kram et al. 2017; Behringer et al. 2024). Our longitudinal model and the inferred fitness seascape thus link short-term competitive fitness effects, with their associated trade-offs and constraints, to long-term adaptive outcomes.

The extended growth curve provides a compelling model of environmental fluctuation because its major phases are well-characterized, reproducible, and biologically interpretable: processing through rapid growth, nutrient depletion with broad physiological remodeling, widespread death and release of complex cell debris, and finally prolonged competition during LTSP. Across this 10-day time series, we identified a large set of confidently non-neutral gene disruptions whose selection rates shift in both magnitude and direction between intervals, and the overall fitness distributions themselves change across phases. These distributional shifts underscore that the growth curve and starvation are not individual selective regimes but rather are temporally structured sequences of environments in which alleles can switch from beneficial to deleterious (or vice versa) as constraints change (Zambrano et al. 1993; Abreu et al. 2024). Additionally, the substantial set of missing library genes—enriched for essential processes like translation, replication, lipid metabolism, and one-carbon metabolism—highlights an orthogonal axis of selection: mutations that are strongly deleterious or lethal in these conditions may be purged before the focal experiment begins. This emphasizes the need to interpret fitness landscapes considering library composition, as we have done by calculating selection rates relative to empirically neutral genes (Wetmore et al. 2015; Price et al. 2018). Another key consequence of this temporal structure is that fitness early during growth phase can disproportionately shape cumulative outcomes over the full 10-day experiment, where early selection may act as a filter on which genetic strategies are available in later phases, consistent with previous studies in similar systems (Katz et al. 2021; Behringer et al. 2024; Patton et al. 2025).

Beyond identifying overall fitness effects, time-resolved inference also makes it possible to identify coherent temporal strategies. Clustering of interval-specific selection-rate trajectories revealed groups of mutants that share characteristic fitness patterns over the growth curve and are enriched for distinct functional modules. Several functional classes with strong fitness effects illustrate how temporally structured environments can produce similar fitness trajectories among mechanistically different mutants. Efflux-related loci highlight redundancy at the level of individual pumps contrasted with the broad importance of TolC as a shared outer-membrane channel (Zgurskaya and Nikaido 2000;

Tikhonova et al. 2011). Biofilm-associated systems contribute in strongly phase-specific ways, with adhesion and aggregation mechanisms diverging between early growth and LTSP (Danese 2000; Chapman et al. 2002; Woude 2008). Finally, regulatory perturbations such as disruption of *cytR* produce some of the strongest effects detected in this experiment; however, some of the factors with the largest regulons—like sigma factors *rpoD* and *rpoS*—were not present in our library. Collectively, these patterns suggest that extended culture selects for various physiological programs that can be reached through multiple genetic routes. These alternative strategies could be responsible for repeated emergence of distinct coexisting ecotypes within the same environment observed in long-term evolution experiments under similar starvation or fluctuation regimes (Finkel and Kolter 1999; Kram et al. 2017; Ratib et al. 2021; Behringer et al. 2022).

Enabled by accurate genome-wide fitness measurements, we further showed that diverse gene-level and pathway-level fitness trajectories can be organized by a low-dimensional latent structure. We formalized this by fitting a one-dimensional geometric “seascape” model in which each mutant is assigned a coordinate on a latent phenotypic axis, and each interval has its own quadratic fitness surface along that axis. The fitted seascape recapitulates the antagonistic selection pressures inferred from interval-specific analyses and previous empirical studies: high fitness during death phase (1→4 d) is inversely correlated with high fitness during early growth and, to a lesser extent, survival during LTSP (Maharjan and Ferenci 2013). The decreasing curvatures over time suggest that the growth/survival trade-off is strongest from exponential growth through death phase; future studies could increase temporal sampling resolution from the onset of death phase into LTSP to determine if the period from 4 d to 10 d has distinct selective pressures with their own trade-offs. Integrating interval-specific landscapes yields a unimodal cumulative fitness peak over 10 days of culture, illustrating how temporally structured environments introduce constraints as the milieu changes, which limits the accessible set of high-fitness strategies available. Importantly, the latent phenotypic axis coordinate provides a common ordering of mutants that links mechanistically disparate loci to a shared ecological interpretation; this organizing axis is a resource that can be used as a predictor or covariate of other assays of knockout mutants.

We further interpreted this latent axis by mapping a large, published compendium of RB-TnSeq experiments onto the seascape. We have shown that genes at low negative phenotypic coordinates tend to have weak, near-neutral effects across growth conditions, whereas genes at high positive phenotypic coordinates increasingly diverge in fitness, producing a gradient from very high fitness in the most stressful conditions to very low fitness in conditions with single carbon or nitrogen sources, which require a diverse metabolic capacity to grow. This latent axis thus appears to be a continuum from environmentally robust but weak-effect alleles to antagonistically pleiotropic alleles that are specialized for stress at the expense of growth. This trade-off at high-phenotypic-coordinate alleles aligns with established growth-stress trade-off paradigms, particularly the trade-off between self-preservation and nutritional competence (SPANC) (Ferenci 2005; Phan and Ferenci 2013).

Finally, we have shown an association between our latent phenotypic axis and transposon fitness effects and the genetic targets of adaptation in an independent long-term evolution experiment under related starvation conditions. Across repeated feast/famine evolution, genes whose disruption had large-magnitude fitness effects—whether strongly beneficial or strongly deleterious—were disproportionately likely to acquire mutations in the evolution experiment, and mutations in these genes arose earlier in the experiment than those in genes with smaller effects. Moreover, transposon insertions in genes toward the negative end of the latent axis (aligned with both the optima for early growth and LTSP and with the integrated 10-day optimum) arose earlier in the long-term experiment than in genes at the positive end of the axis. These results underscore the utility of our approach in predicting evolutionary outcomes: transposon mutants with large effects, whether positive or negative, possibly signal genes with the potential to produce any large effect via mutation and thus shift genotypes into previously inaccessible regions of phenotypic space where new adaptive paths become available; however, our seascape model takes into account the relative magnitudes of temporal trade-offs and is therefore more narrowly predictive of the fitness benefits of transposon insertions under this particular selective regime. Thus, this dual experimental–modeling approach indicates that transposon mutant fitness can serve as a meaningful indicator of loci that repeatedly respond to and reshape the fitness landscape under strong, structured selection.

Together, this work highlights the utility of longitudinal RB-TnSeq, paired with principled Bayesian inference, as a general platform for mapping selective dynamics in temporally structured environments. By quantifying time-dependent fitness effects, compressing complex gene-by-time patterns into an interpretable latent axis, and testing how regulatory structure and evolutionary outcomes interact with that axis, we provide a framework for connecting mechanistic gene functions to systems-level constraints and long-term adaptation. While our study focused on starvation in *E. coli*, this same strategy is broadly extensible to other time scales, stress sequences, and community contexts, enabling a broader empirical and theoretical synthesis of how fluctuating environments shape fitness landscapes and evolutionary predictability.

## METHODS

### Strain and culture conditions

All experiments were conducted using Escherichia coli K-12 strains. The wild-type strain was *E. coli* K-12 substr. MG1655 (Strain PFM2 from Lee *et al*. (2012)). Transposon mutant fitness was measured using the randomly barcoded transposon library KEIO_ML9, which was previously constructed by random insertion of barcoded Tn5 transposons into BW25113, the parent strain of the Keio deletion collection (Baba et al. 2006; Wetmore et al. 2015). The original transposon library contained 3728 unique transposon insertions with a median number of 16 unique insertions per gene (Wetmore et al. 2015). Unless otherwise specified, strains were cultured in LB-Miller medium (10 g/L tryptone, 5 g/L yeast extract, 10 g/L NaCl) at 37 °C with orbital shaking at 175 rpm. Cultures were grown in 16 x 100 mm glass culture tubes positioned upright in the shaking incubator.

### Longitudinal RB-TnSeq competition assay

To quantify fitness dynamics across the extended growth curve, we performed longitudinal RB-TnSeq assays spanning 10 days of culture. Frozen aliquots of the KEIO_ML9 library were thawed on ice and inoculated into 50 mL LB with kanamycin (30 ug/mL) in a 250-mL flask, then cultured to mid-exponential phase (OD_600_ ≍ 0.5; V-1200 Spectrophotometer, VWR International, Radnor, PA, USA). In parallel, wild-type MG1655 cultures were grown under the same conditions without antibiotics, starting from an overnight culture diluted to OD_600_ = 0.05.

Both library and wild-type cultures were washed twice in PBS and normalized to OD_600_ = 1.0. Co-cultures were initiated by mixing 100 μL of KEIO_ML9 with 100 μL of wild-type cells into 10 mL of fresh LB medium in a culture tube. Parallel monocultures of KEIO_ML9 alone were also established under identical conditions. Three 1-mL aliquots of the KEIO_ML9 library were pelleted and frozen for time 0 samples.

Because upright test tube cultures develop spatially structured microenvironments that are disrupted by sampling, we employed a pseudo-longitudinal design (Behringer et al. 2018). Independent replicate cultures were established for each sampling time point using equal volume inocula from the same starting cultures. For each time point, 3 biological replicate cultures were grown each for the KEIO_ML9 library alone and the KEIO_ML9 library mixed with wild-type, resulting in 6 biological replicates per time point that were ultimately pooled after bioinformatic processing. Samples were collected immediately after inoculation (day 0) and after 1, 4, and 10 days of incubation. For day 0, three 1-mL aliquots of the KEIO_ML9 library were collected prior to incubation. At each subsequent time point, 1 mL samples were harvested, pelleted by centrifugation, and stored at −80 °C for until DNA extraction.

### DNA isolation, barcode amplification, and sequencing

DNA extraction, barcode amplification, and DNA sequencing were performed similarly to previously published uses of the RB-TnSeq library (Wetmore et al. 2015; Price et al. 2018). Genomic DNA was isolated from frozen pellets using the Omega Bio-tek E.Z.N.A Bacterial DNA Kit (Product No. D3350) following the manufacturer’s protocol, including the optional mechanical lysis with glass beads (Omega Bio-tek, Norcross, GA, USA). Transposon barcodes were then amplified using the BarSeq98 protocol as previously described (Wetmore et al. 2015). Each sample was amplified using a sample-specific indexed forward primer (ITXX) and a common reverse primer (P1). PCR amplification followed the BarSeq98 protocol, with an initial denaturation at 98 °C for 4 min, followed by 25 cycles at 98 °C for 30 s, 55 °C for 30 s, and 72 °C for 30 s, with a final extension step of 72 °C for 5 min. Samples were sequenced together on an Illumina NovaSeq6000 instrument at the Vanderbilt VANTAGE genomics core facilities using paired-end 150-bp chemistry; only read 1 was used for downstream analysis. Before barcode counting, reads were trimmed using Trimmomatic v. 0.39 single-end mode to remove Illumina adapter sequences and crop reads to 75 base pairs to reduce file sizes (Bolger et al. 2014).

### Barcode processing and normalization

Barcode identification, initial filtering, and counting were performed using previously published scripts from the RB-TnSeq pipeline (Wetmore et al. 2015). Downstream analyses were performed using custom R scripts developed for this study.

Barcodes were first identified from reads and counted using script MultiCodes.pl, after which barcode locations were mapped to Tn insertion locations using the pool definition file and combineBarSeq.pl (Wetmore et al. 2015). We then used the “poolcount” file output from combineBarSeq.pl in downstream analysis. Barcodes were filtered to remove any barcode with fewer than three total counts across all day 0 samples. To mitigate positional sequencing biases, read counts were corrected by subtracting a sliding median calculated across 251-barcode windows. Barcode counts were then normalized by log-ratio transformation relative to an empirically defined neutral gene set. Log-ratio transformation was done using R package ALDEx2, using the clr() function with neutral genes input as the “denom” option (**Supplemental Text 1**) (Fernandes et al. 2013; Gloor et al. 2017). Neural genes were identified based on a large published RB-TnSeq compendium as genes whose disruption exhibited minimal fitness variation across diverse environmental conditions (**Table S1**) (Price et al. 2018).

### Calculation and Bayesian inference of selection rates

Selection rates were calculated from normalized barcode abundances as the change in log-ratio abundance between adjacent time points divided by the elapsed time in days. Selection rates were calculated separately for each barcode across three intervals: day 0→1 (*s*_0→1_), day 1→4 (*s*_1→4_), day 4→10 (*s*_4→10_). This sliding-window approach enables time-resolved inference of mutant fitness across distinct physiological phases of the extended growth curve.

Selection rates were then modeled using Bayesian multilevel piecewise linear regression. Each gene was treated as a group-level effect, with separate slopes estimated for days 1-4 and 4-10, and intercepts reflecting the day 1 selection rate. The model structure incorporated an LKJ prior on the correlation between the intercept and slopes and regularizing Student-t priors on all hyperparameters to stabilize inference given the relatively small number of barcodes per gene (Extended methods in **Supplemental Text 1**). Population-level priors were parameterized based on reanalysis of published RB-TnSeq data (Price et al. 2018).

Posterior distributions were first approximated using the Pathfinder variational inference algorithm implemented in CmdStanR, accessed through the R package brms (Hoffman and Gelman 2011; Bürkner 2017; Zhang et al. 2022; Gabry et al. 2025). This approximation allowed efficient initialization of full Hamiltonian Monte Carlo (HMC) sampling, greatly reducing computation time (Neal 2011). Final HMC sampling was performed with draws saved for downstream analysis. Model convergence and effective sample sizes were assessed to ensure accurate posterior estimation. This hierarchical Bayesian approach provided shrinkage of noisy estimates toward neutrality while preserving strong fitness effects.

### Summary statistics and functional analysis

Posterior probability distributions of the expected values were generated for each gene’s selection rates. From these estimates, cumulative log relative abundance was calculated as the integral of the step function of selection rate over the full 10-day culture period (**Supplemental Text 1**). For each gene, we summarized fitness effects using both point estimates (posterior medians) and probability thresholds (e.g., probability > 90% that fitness deviates from neutrality). Genes were clustered based on selection rate trajectories using principal component analysis and k-means clustering (k = 4). Function enrichment analyses (GO, KEGG, EcoCyc pathway annotations) were performed within each cluster to identify biologically coherent groups (Ashburner et al. 2000; Kanehisa and Goto 2000; Kanehisa et al. 2025; Karp et al. 2025; The Gene Ontology Consortium 2026). Functional enrichment was done in R using package clusterProfiler and the latest Bioconductor release of the *E. coli* K-12 genome annotation (v. 3.22.0) (Yu et al. 2012; Wu et al. 2021; Carlson 2024).

### Benchmarking simulation

Precision and accuracy of predicted fitness values was measured by analyzing simulated genes with noise from multiple sources. Genes were simulated with latent knockout selection rate effects. As in our biological system, the selection rate and abundance of each gene was simulated forward through three time points, with selection rates of each gene correlating across time. Noise was introduced in three ways: selection rates of each barcode were sampled from a normal distribution around the latent gene selection rate (called intragenic fitness variation); noise was added to each observation to simulate measurement error with no directional bias (called measurement error); and a directional bias was added to all observations for each replicate sample simulated (called batch effect). Data were simulated with every combination of these three noise parameters generated from normal distributions with standard deviations of 0, 0.1, or 1 with units of days^−1^, as in selection rate. For each noise combination, selection rate over time was estimated using a statistical inference pipeline designed for RB-TnSeq data (BarSeq, rewritten in R from equations in Wetmore et al. 2015) or with our longitudinal inference model. For each condition, 1000 genes were simulated and root mean square error (RMSE) and coverage probability, measured as the proportion of true selection rates within inferred 90% confidence intervals, were calculated, with nonparametric bootstrapping used to estimate confidence around these two statistics. To better parse the effects of various sources of noise, RMSE and coverage were then modeled as a correlated multivariate generalized linear model with main effects and interactions of the three noise parameters as predictors, hierarchically grouped by inference method. The effects of each inference method on RMSE and coverage were then estimated conditioned on the levels of each noise parameter using conditional_effects() in brms.

### Geometric seascape model of time-dependent fitness

To summarize structured changes in mutant library fitness across the three measured intervals, we fit a one-dimensional Fisher’s geometric seascape model to the interval-specific gene selection-rate estimates from our longitudinal RB-TnSeq model. Briefly, the seascape assigns each gene a latent coordinate 𝑧_g_ on a one-dimensional phenotypic axis and estimates an interval-specific quadratic fitness surface characterized by an optimum (𝜃_𝑑_) and curvature (𝜅_𝑑_):

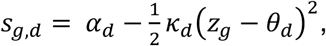

where α_I_ is the maximum selection rate at the optimum. To encode temporal dependence without enforcing a linear or directional trend, interval optima were regularized with a discretized Ornstein-Uhlenbeck process linking adjacent intervals (Uhlenbeck and Ornstein 1930; Hansen 1997). The model was fit in Stan, using posterior summaries from the longitudinal model as inputs (carrying forward uncertainty), and posterior draws were used to compute gene coordinates, interval-specific landscapes, and the integrated 10-day landscape. Full likelihood, priors, and parameterization are provided in Supplemental Text 1.

This model was fit jointly across all genes and intervals using Bayesian inference in Stan using R package *brms* (Bürkner 2017; Gabry et al. 2025; Stan Development Team 2025). Observed gene-level selection rate posterior estimates from the longitudinal RB-TnSeq model were treated as noisy observations of the latent seascape-predicted selection rates. Weakly informative priors were placed on all parameters to regularize inference while allowing flexibility in landscape shape and temporal dynamics.

Cumulative fitness across the full 10-day experiment was calculated by summing interval-specific fitness functions weighted by interval duration, producing an integrated seascape with a single cumulative optimum. All downstream analyses of latent phenotypic coordinates used full posterior distributions to propagate uncertainty.

## Data and code availability

Sequencing data for RB-TnSeq experiments can be downloaded from SRA, BioProject PRJNA1422734. Code, count tables, and files required to reproduce this analysis are found at https://github.com/BehringerLab/Longitudinal_RBTnSeq_Paper.

## Acknowledgments

We would like to thank J.B.M. for providing the KEIO_ML9 RB-TnSeq library. We would like to thank Vanderbilt Technologies for Advanced Genomics (VANTAGE) for DNA sequencing and high-performance computing resources provided by the Advanced Computing Center for Research and Education (ACCRE) at Vanderbilt University. This work was supported by Army Research Office Grant W911NF-21-1-0161 (M.G.B.) and National Institutes of Health Grant R35GM150625 (M.G.B.).

## Author Contributions

C.J.S. and M.G.B.: conceptualization, investigation, data visualization, writing, and editing. C.J.S.: statistical analysis, including development of the longitudinal model and seascape model. M.G.B.: funding acquisition, resources, supervision.

