## Supplemental Materials for "Bayesian analysis of longitudinal RB-TnSeq resolves the fitness seascape in fluctuating environments"

Text S1. Extended Methods: Model specifications for longitudinal RB-TnSeq inference and the geometric seascape

Figure S1. Accuracy and uncertainty of our longitudinal RB-TnSeq model compared to the standard method.

Figure S2. Variability of selection rates within genes, between replicates, and by experimental batch.

Figure S3. Enriched functions of transposon mutants not detected in the transposon library.

Figure S4. Variability versus effect size of predicted selection rate.

Figure S5. Selection rate distributions and skewness by time interval.

Figure S6. Clustering scores for determining *k*-means clusters of non-zero fitness effects.

Figure S7. Mutants with strong fitness effects display potential cumulative network fitness effects and time-dependent selection.

Figure S8. The *cytR* regulon contains more deleterious mutants than expected under size-matched null gene sets.

Figure S9. Fitness across diverse growth conditions mapped to the seascape coordinates.

Figure S10. Seascape coordinate predicts allele arrival time across certain mutation classes.

Figure S11. Locus-specific mutation frequency trajectories across repeatedly mutated genes in feast/famine evolution.

**Additional Files include Supplemental Tables and Dataset S1:**

Table S1: Neutral loci from previous RB-TnSeq experiments.

Table S2: Genes present in the original RB-TnSeq ML9 library but not detected in our analysis.

Table S3: Number of confidently non-zero transposon mutants by time interval with a strict probability cutoff of 97.5%.

Table S4: Functional, pathway, and gene ontology enrichment of transposon mutants with non-neutral fitness effects, separated by fitness trajectory cluster.

Table S4: RB-TnSeq experiments reanalyzed from the Fitness Browser.

Table S5: RB-TnSeq experimental conditions clustered by similarity of fitness profiles.

Table S6: Clusters and broad descriptions of Fitness Browser RB-TnSeq experimental conditions.

Dataset S1. Selection rates of transposon insertion mutants estimated with a longitudinal Bayesian hierarchical model.

**Supplemental Text S1**

### Extended Methods: Model specifications for longitudinal RB-TnSeq inference and the geometric seascape

This supplement describes in detail the Bayesian models used in this study for the purposes of reproducibility and future iterations. Code corresponding to these models can be found on the Github repository for this paper. All analyses were done using R v. 4.4.2, brms v. 2.23.0, cmdstanr v. 0.9.0.9000, CmdStan v. 2.36.0, and Stan v. 2.37.

#### Data and notation

The KEIO_ML9 RB-TnSeq library was sampled at four time points (day 0, 1, 4, 10), which define three adjacent intervals: 0→1, 1→4, and 4→10. Each gene in the library has on average 16 unique transposon insertion mutants, each with a unique barcode. Let $c_{b,t}$ be the sequencing count for barcode $b$ at time $t$. Counts were normalized using a centered log-ratio transformation relative to an empirically neural gene set (implemented in ALDEx2 using the neutral set as the denominator), giving the normalized log-ratio abundance $y_{b,t}$.

For each interval $t \to t'$, the barcode-level selection rate on a per-day scale is defined as

$$r_{b,(t \to t')} = \frac{y_{b,t'} - y_{b,t}}{t' - t}.$$

For conciseness, the interval will be referred to by its ending day $d \in \{1,4,10\}$, so $d = 1$ denotes $0 \to 1$, $d = 4$ denotes $1 \to 4$, and $d = 10$ denotes $4 \to 10$. These $r_{b,d}$ values are then used as the independent variable input to our longitudinal model.

#### Multilevel model for gene-by-interval selection rates

Our longitudinal selection rate model uses a piecewise linear regression formula with hierarchical modeling of gene-specific mean selection rate inferred from barcode-level fitness trajectories. For each observed log-ratio-transformed barcode-level selection rate $r_{b,d}$, specific to gene $g[b]$ and interval $d$, we use a robust Student-*t* likelihood,

$$r_{b,d} \sim \text{Student-}t\big(v_d, \mu_{g[b]}(d), \sigma_d\big),$$

where both the residual scale $\sigma_d$ and the degrees of freedom $v_d$ vary with interval $d$. This is parameterized as

$$v_d = \beta_0^{v} + \beta_1^{v} d$$

with priors

$$\beta_0^{v} \sim \text{Normal}(4, 0.8),$$

$$\beta_1^{v} \sim \text{Normal}(0, 0.05),$$

and

$$\log \sigma_d = \beta_0^{\sigma} + \beta_1^{\sigma} d$$

with priors

$$\beta_0^{\sigma} \sim \text{Normal}(-2, 0.2),$$

$$\beta_1^{\sigma} \sim \text{Normal}(-0.5, 0.1).$$

The gene-level mean $\mu_g(d)$ is modeled as a continuous piecewise-linear function with a knot at day 4:

$$\mu_g(d) = \beta_{0g} + \beta_{1g} d + \beta_{2g}(d-4)\mathbb{I}(d \geq 4).$$

This produces two linear fits continuous at the knot at day 4, where $\beta_{1g}$ is the slope from $d = 1$ to $d = 4$, and the slope from $d = 4$ to $d = 10$ is $\beta_{1g} + \beta_{2g}$.

Gene-specific parameters are partially pooled using a correlated multivariate normal prior:

$$\begin{pmatrix} \beta_{0g} \\ \beta_{1g} \\ \beta_{2g} \end{pmatrix} \sim \text{MVNormal}\left( \begin{pmatrix} \bar{\beta}_0 \\ \bar{\beta}_1 \\ \bar{\beta}_2 \end{pmatrix}, \boldsymbol{\Sigma} \right), \qquad \boldsymbol{\Sigma} = \text{diag}(\tau_0, \tau_1, \tau_2)\, \mathbf{R}\, \text{diag}(\tau_0, \tau_1, \tau_2),$$

where $\mathbf{R}$ is given an LKJ prior. This correlation structure shrinks genes with few or noisy barcodes toward the population mean trajectory while allowing genes with consistent barcodes or very strong effects to pull away from the mean.

Since population-level mean selection rates are expected *a priori* and by design to be close to zero, moderately informative priors were used to regularize noisy estimates while allowing for rare large effects:

$$\bar{\beta}_0 \sim \text{Normal}(0, 0.01)$$

$$\bar{\beta}_1 \sim \text{Normal}(0, 0.005)$$

$$\bar{\beta}_2 \sim \text{Normal}(0, 0.005).$$

Group-level standard deviations are given priors

$$\tau_k \sim \text{Student-}t(9,0,0.005)$$

for $k \in \{0,1,2\}$ with $\tau_k > 0$, and

$$\mathbf{R} \sim \text{LKJ}(4).$$

Since this model has thousands of parameters that make full Monte Carlo sampling difficult to initialize, we first used the Pathfinder algorithm to find a stable region of parameter space and produce an approximate posterior, which was then used to initialize a full Hamiltonian Monte Carlo sampling solution. Convergence and sampling pathologies were assessed with standard diagnostics ($\hat{R}$, effective sample size, and divergent transitions).

Posterior draws of $\mu_g(d)$ at $d \in \{1, 4, 10\}$ are the reported interval-specific gene selection rates,

$$s_{0\to1,g} = \mu_g(1), \qquad s_{1\to4,g} = \mu_g(4), \qquad s_{4\to10,g} = \mu_g(10).$$

From these, the cumulative expected change in log-ratio abundance across $0 \to 10$ days is the time-weighted sum,

$$\Delta_g = 1 \cdot s_{0\to1,g} + 3 \cdot s_{1\to4,g} + 6 \cdot s_{4\to10,g},$$

and the net per-day selection rate across the full experiment is

$$s_{net,g} = \frac{\Delta_g}{10}.$$

**A Fisher's geometric seascape model with a moving OU optimum**

Our seascape model compresses gene-by-interval selection rate trajectories in a latent one-dimensional coordinate $z_g$ for each gene by jointly estimating a single-optimum fitness landscape for each interval. The gene-level selection rate estimates from our longitudinal model were used to fit the seascape model, carrying forward uncertainty from posterior predictions. Let $\tilde{s}_{g,d}$ be the gene-level estimate for gene $g$ in interval $d$, and let $\tilde{\sigma}_{g,d}$ be the posterior standard deviation. We treat $\tilde{s}_{g,d}$ as an observation of gene latent mean selection rate $m_{g,d}$ with a measured standard error plus additional residual deviation estimated by the model:

$$\tilde{s}_{g,d} \sim \text{Normal}\left(m_{g,d}, \sqrt{\tilde{\sigma}_{g,d}^2 + \sigma^2}\right).$$

The seascape mean function is a quadratic surface determined by distance from the optimum:

$$m_{g,d} = \alpha_d - \tfrac{1}{2}\kappa_d\left(z_g - \theta_d\right)^2,$$

where $\theta_d$ is the interval-specific optimum, $\alpha_d$ is the maximum selection rate at the optimum, and $\kappa_d > 0$ is the curvature, proportional to the strength of stabilizing selection around the optimum. The optimum location, curvature, and maximum selection rate are estimated separately by interval.

To encode temporal dependence in the selective regime, the optima $\{\theta_1, \theta_4, \theta_{10}\}$ were modeled by a discretized Ornstein-Uhlenbeck (OU) process around an estimated long-run mean $\mu$. Between each interval transition, the next optimum is influenced by $\mu$ and a Gaussian innovation term $\epsilon$

$$\theta_{t'} = \mu + (1 - \rho)(\theta_t - \mu) + \phi\epsilon$$

where $\rho \in (0, 1)$ determines the strength of reversion to the mean, and $\phi$ determines the strength of innovation. We parameterized $\rho$ on a logit scale and innovation strength as

$$\phi = \omega\sqrt{\rho(2 - \rho)}$$

for sampling stability. $\rho$ and $\phi$ were estimated separately for the $1 \to 4$ and $4 \to 10$ transitions, with $\omega$ shared between the two transitions but with an added offset in $\omega$ estimated for the $4 \to 10$ transition

$$\phi_{4\to10} = \omega\, \Delta_\omega \sqrt{\rho_{4\to10}(2 - \rho_{4\to10})}$$

with a weak prior on $\Delta_\omega$

$$\log \Delta_\omega \sim \text{Normal}(0, 0.35).$$

The long-run mean is constrained positive with prior

$$\log \mu \sim \text{Normal}(0, 0.15),$$

and the initial optimum is set below $\mu$ as

$$\theta_1 = \mu - e^{\Delta_{\theta 0}}.$$

The optima at days 4 and 10 are then propagated forward by OU transitions:

$$\theta_4 = \mu + (1 - \rho_{1\to4})(\theta_1 - \mu) + \phi_{1\to4}\epsilon_{1\to4}$$

$$\theta_{10} = \mu + (1 - \rho_{4\to10})(\theta_4 - \mu) + \phi_{4\to10}\epsilon_{4\to10}$$

with priors

$$\text{logit}(\rho) \sim \text{Normal}(0, 0.4),$$

$$\epsilon_{1\to4} \sim \text{Normal}(0, 0.7), \qquad \epsilon_{1\to4} \geq 0,$$

$$\epsilon_{4\to10} \sim \text{Normal}(0, 1)$$

Arbitrary constraints were added to some parameters to prevent symmetrical solutions and improve identifiability. First, $z_g$ is partially pooled per gene but constrained to have a population mean of 0 and standard deviation of 1. Next, along with the constraint that $0 < \theta_1 < \mu$, the initial innovation step $\epsilon_{1\to4}$ is constrained to be positive. This parameterization ensures that the relative orientations of the latent axis, $z_g$ ordering, optima, and OU steps are reproducible across model runs.

Given these constraints, priors on all other parameters were weakly informative, encoding a descending maximum selection rate over time (as qualitatively observable from raw data and our longitudinal model) and a larger curvature in the $1 \to 4$ interval (due to exponential growth phase allowing larger relative fitness changes compared to later growth states):

$$\alpha_1 \sim \text{Normal}(1.5, 0.5),$$

$$\alpha_4 \sim \text{Normal}(1.0, 0.3),$$

$$\alpha_{10} \sim \text{Normal}(0.3, 0.1),$$

$$\log \kappa_1 \sim \text{Normal}(-0.5, 0.3),$$

$$\log \kappa_4 \sim \text{Normal}(-1.5, 0.3),$$

$$\log \kappa_{10} \sim \text{Normal}(-1.5, 0.3)$$

Computation was done using Stan's HMC sampler algorithm using 4 chains with 1000 warmup iterations and 2000 sampling iterations per chain.

Cumulative summaries across the full experiment were computed across all posterior draws to calculate the integrated landscape from all three intervals. Each interval was weighted by its duration $\Delta t_d \in \{1, 3, 6\}$ so that the cumulative expected selection rate inferred from the seascape is

$$\tilde{S}_g^{0\to10} = \sum_{d\in\{1,4,10\}} \Delta t_d \left[\alpha_d - \frac{1}{2}\kappa_d\left(z_g - \theta_d\right)^2\right].$$

The cumulative optimum $z^*$ can then be computed as the coordinate $z$ that maximizes the cumulative selection rate as estimated by the seascape model:

$$z^* = \frac{\sum_d \Delta \mathrm{t_d} \kappa_d \theta_d}{\sum_d \Delta t_d \kappa_d}.$$

All parameters derived from the seascape model including $z^*$ are calculated from posterior draws to propagate uncertainty.

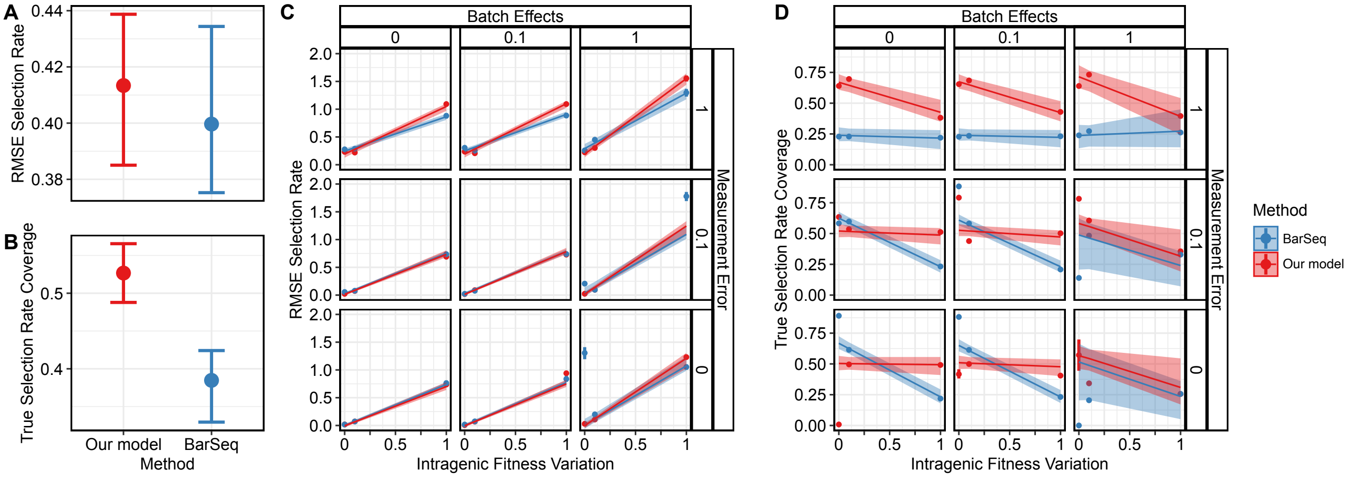


**Figure S1. Accuracy and uncertainty of our longitudinal RB-TnSeq model compared to the standard method.** Simulated gene fitness values were inferred using our longitudinal Bayesian model and a standard BarSeq-style estimator. Performance was quantified by RMSE and probability coverage of true values, then summarized by modeling statistics as functions of simulated noise parameters. **(A)** RMSE and **(B)** coverage of our model and BarSeq conditioned on average noise parameter values. Points are medians with bars for 95% CI. **(C)** RMSE and **(D)** coverage across increasing noise. Noise parameters were simulated as 0-centered normal distributions with standard deviations shown as the parameter value. Noise was then multiplied (for intragenic variation and measurement error) or added (for batch effects) to the true fitness values. Lines and credible intervals reflect the modeled statistic at each indicated combined noise level. Each point and vertical line is the average estimated statistic with 95% CI estimated by nonparametric bootstrap.

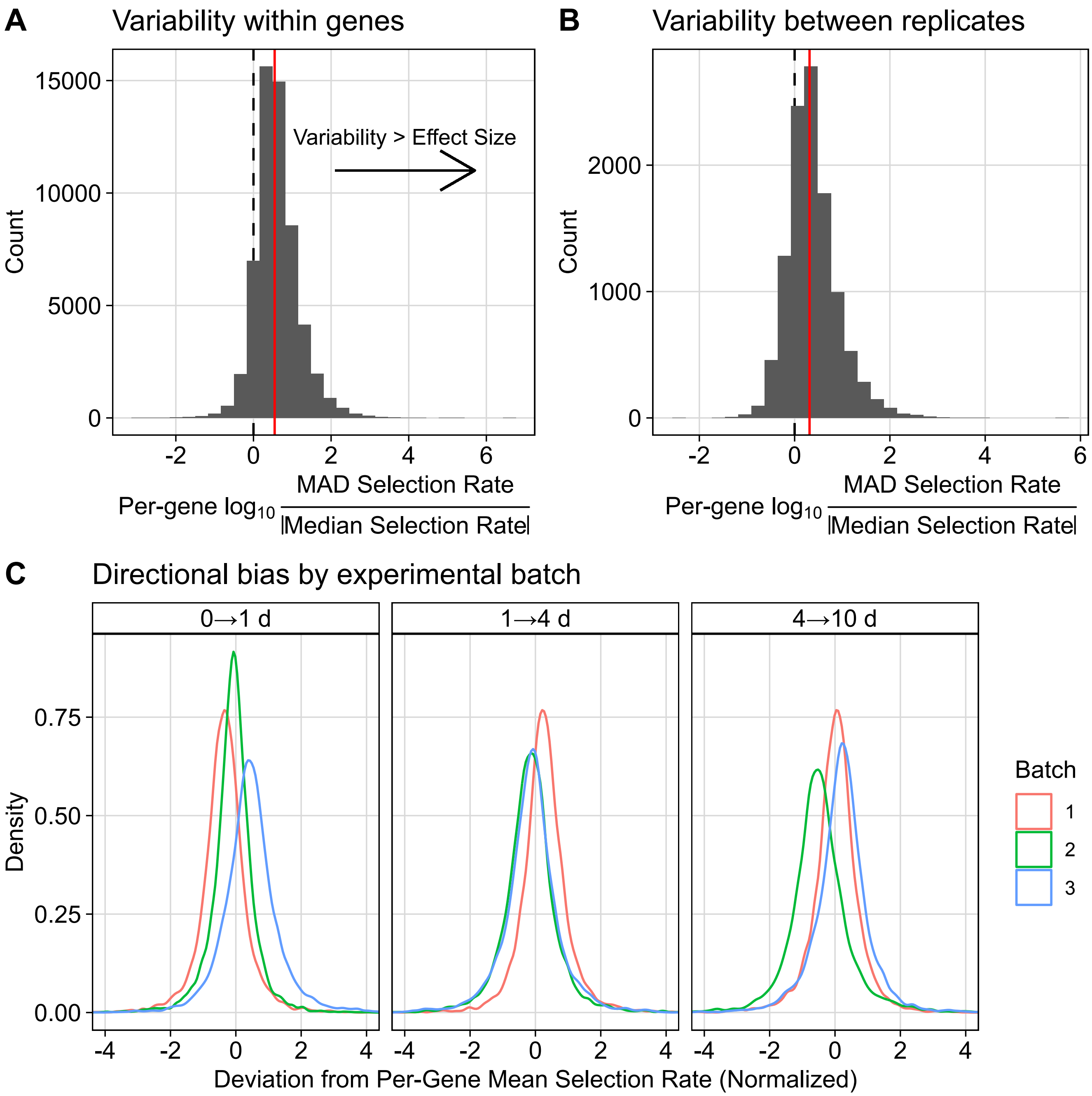


**Figure S2. Variability of selection rates within genes, between replicates, and by experimental batch. (A)** Transposon insertions within genes have much greater variability than the average effect of most genes. **(B)** Selection rate variability between biological replicates is larger than the average selection rate effect of most genes. **(C)** Selection rate was averaged for each gene in each biological sample, after which all selection rates were normalized by day. The difference between the per-gene selection rate of each biological sample was plotted compared to the mean selection rate for that gene across all samples. Differences are separated by experimental batch to show varying directional shifts per day and batch.

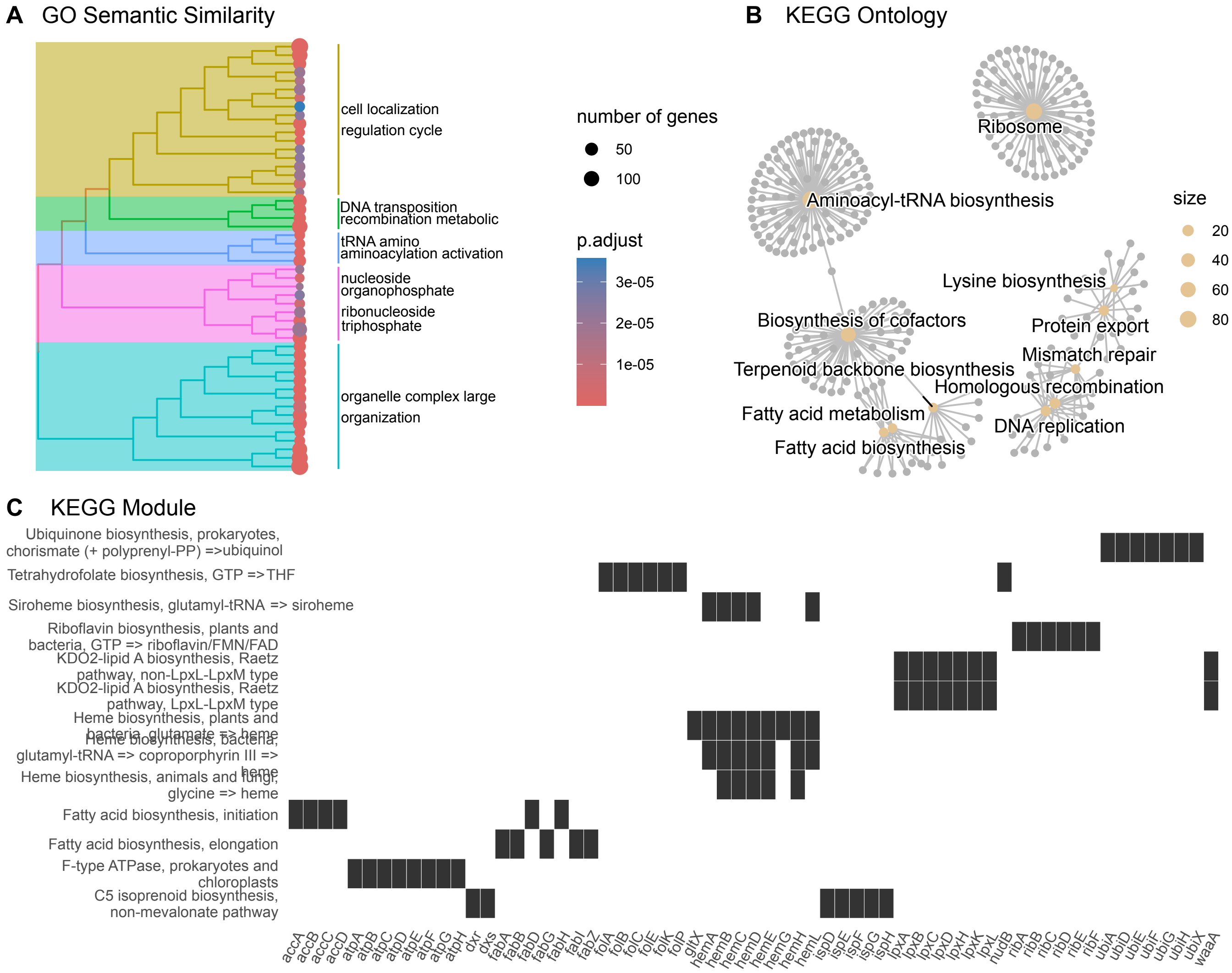


**Figure S3. Enriched functions of transposon mutants not detected in the transposon library. (A)** Semantic similarity of overrepresented gene ontology (GO) terms. Semantic similarity was calculated with Wang's method, and GO terms were clustered by complete linkage. **(B)** Network view of KEGG ontologies. KEGG ontologies are gold nodes with each gene enriched for that ontology linked as a gray node. **(C)** Enriched KEGG modules. For each KEGG module row, genes enriched for that module are indicated by black boxes.

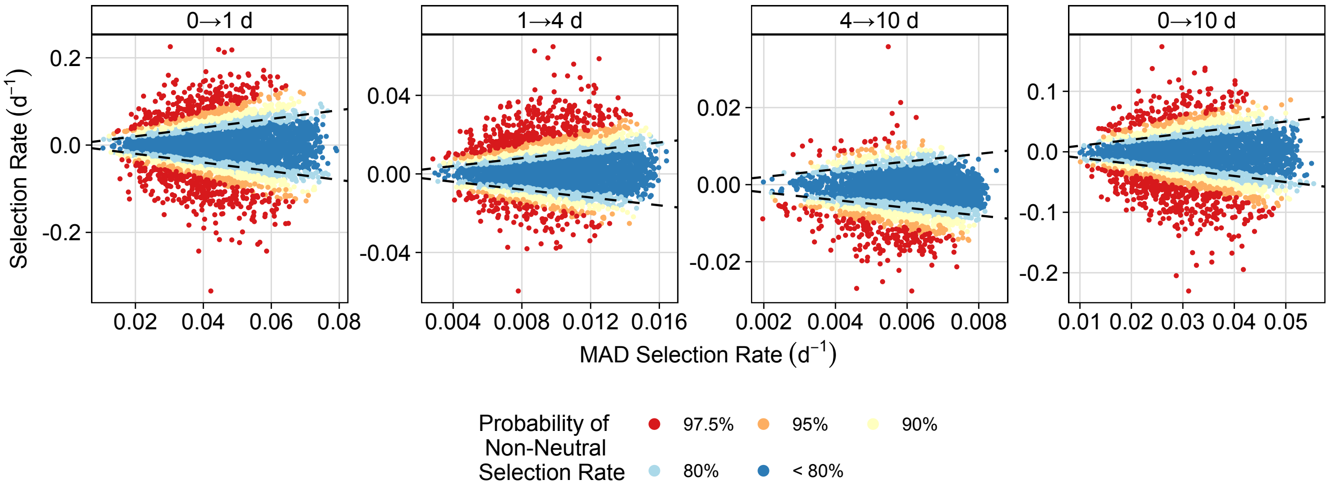


**Figure S4. Variability versus effect size of predicted selection rate.** Each point represents one gene, colored by the posterior probability that the selection rate is above or below 0. Median absolute deviation (MAD) is calculated for each gene's posterior density. Dashed lines indicate the heuristic boundary |median| = MAD, separating genes with effects larger than their posterior dispersion from genes whose inferred effect is equal to or smaller than posterior variability.

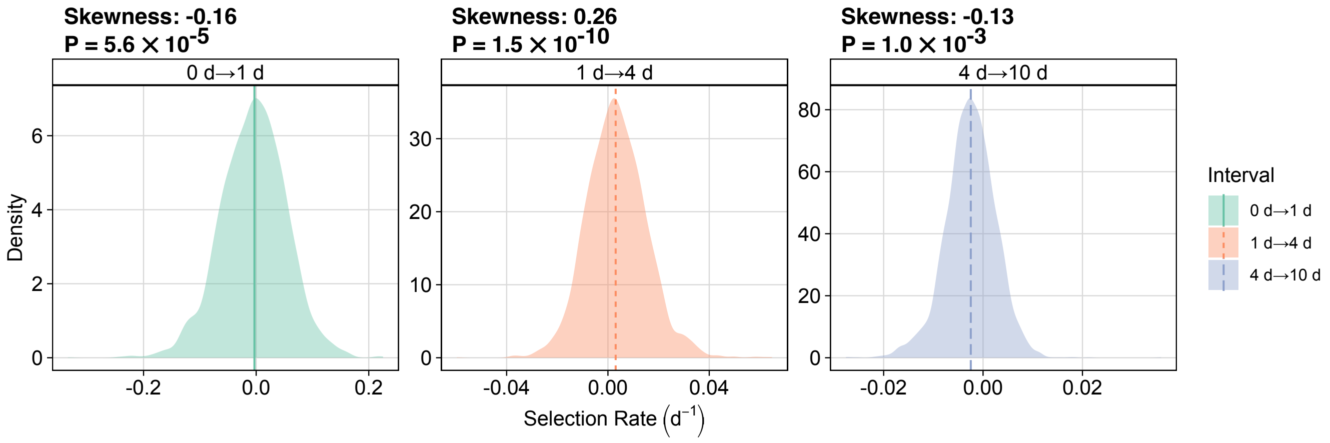


**Figure S5. Selection rate distributions and skewness by time interval.** The posterior medians of selection rate for each gene are shown separated by time interval. Vertical lines indicate the median of each distribution. Sample skewness is indicated above each distribution, with *p*-values from D'Agostino's $K^2$ test for departures from normality (skewness and kurtosis).

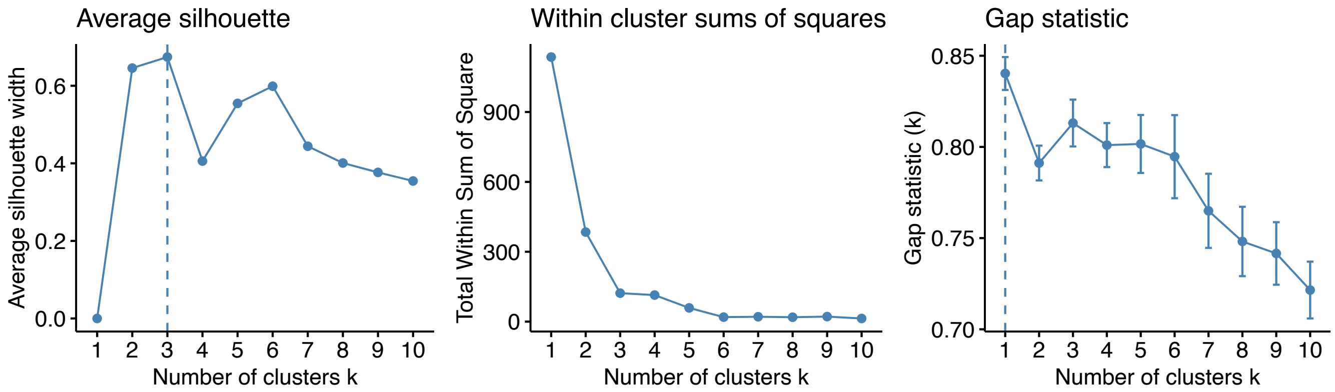


**Figure S6. Clustering scores for determining *k*-means clusters of non-zero fitness effects.** Selection rate estimates for genes with non-zero fitness effects were clustered using *k*-means clustering from *k*=1 to *k*=10. Silhouette width, within-cluster sums of squares, and gap statistic were calculated on all clustering options. *k*=4 was chosen to maximize the number of clusters and biological interpretability while retaining relatively high diagnostic statistic values.

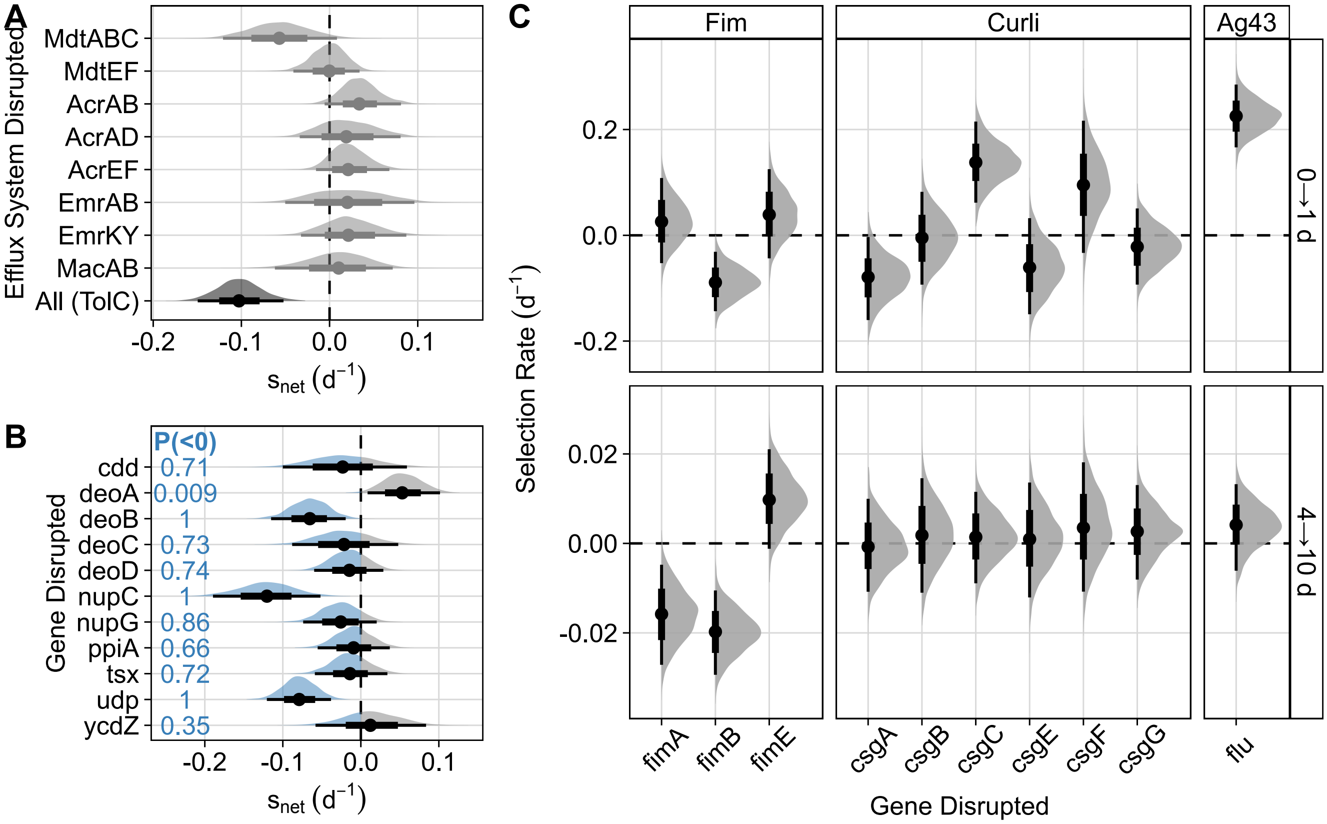


**Figure S7. Mutants with strong fitness effects display potential cumulative network fitness effects and time-dependent selection.** Points and intervals summarize posterior medians with 66% and 95% credible intervals. **(A)** Posterior distributions of $s_{net}$ of efflux-associated mutants. For each efflux system except TolC, the density shown is the combined posterior distribution of each gene listed. For TolC, the density indicates only the posterior distribution of the *tolC* mutant. **(B)** Posterior distributions of $s_{net}$ for genes in the *cytR* regulon. Density below 0 is highlighted in blue, and the probability that $s_{net}$ is less than 0 for each mutant gene is shown. **(C)** Selection rate of biofilm-associated gene mutants across time. Each density and interval indicates the posterior distribution of the Tn mutant indicated.

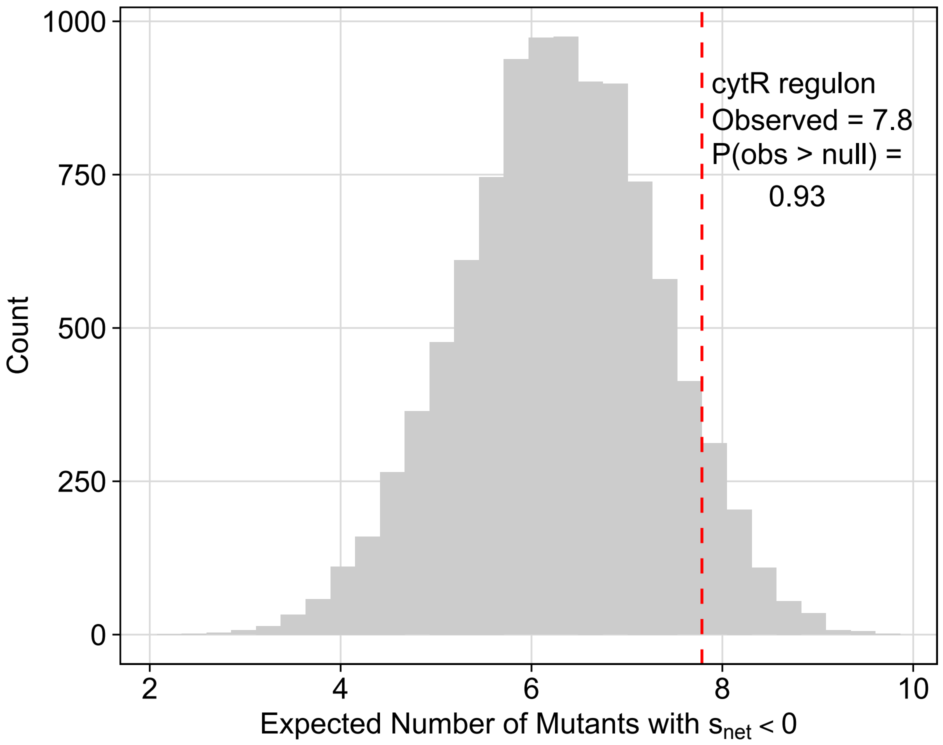


**Figure S8. The *cytR* regulon contains more deleterious mutants than expected under size-matched null gene sets.** For a gene set, the expected number of deleterious mutants was calculated as $\sum_g \Pr\,(s_{net} < 0)$. The null distribution was generated from 10,000 random permutations of size-matched gene sets, and the observed *cytR* regulon expectation is shown relative to the null (dashed red line).

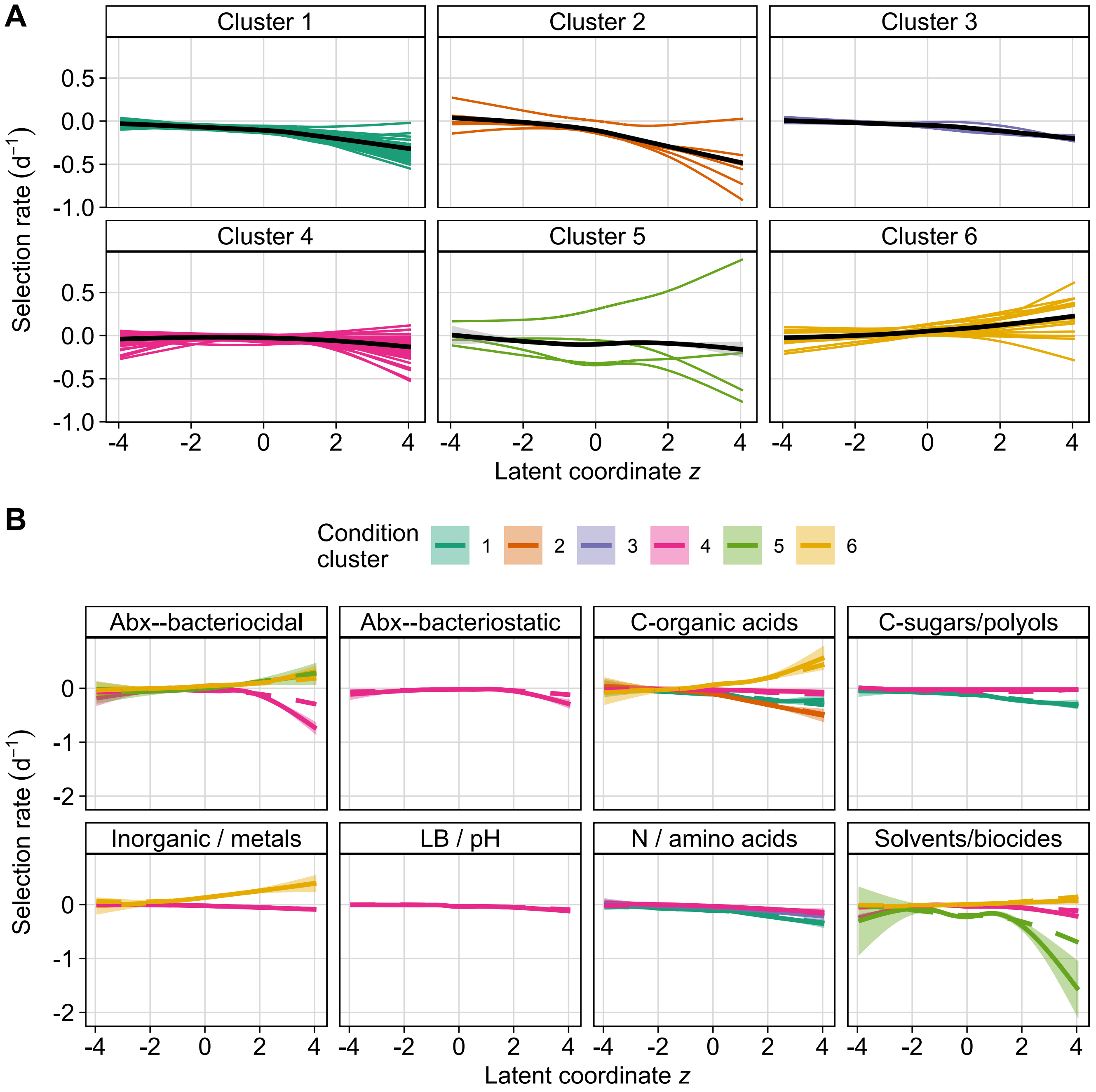


**Figure S9. Fitness across diverse growth conditions mapped to the seascape coordinates. (A)** Condition-specific relationships between fitness and latent *z* coordinates inferred from a generalized additive model. For each condition, the selection rate was modeled as a function of latent coordinate *z*, and predicted selection rates are shown as individual colored lines. Conditions are separated into panels by cluster to visualize the trajectories within and between clusters. Black curves (± 95% CI) are the average predicted trend within each cluster. **(B)** Fitted curves align with raw compendium data within and between clusters and types of conditions. Solid lines (± 95% CI ribbons) are smoothed fits (GAM) to the observed selection rates from the RB-TnSeq compendium as a function of latent coordinate *z*, grouped by condition cluster and separated by the type of conditions. Dashed lines show the corresponding predicted selection rates from the same model as panel A.

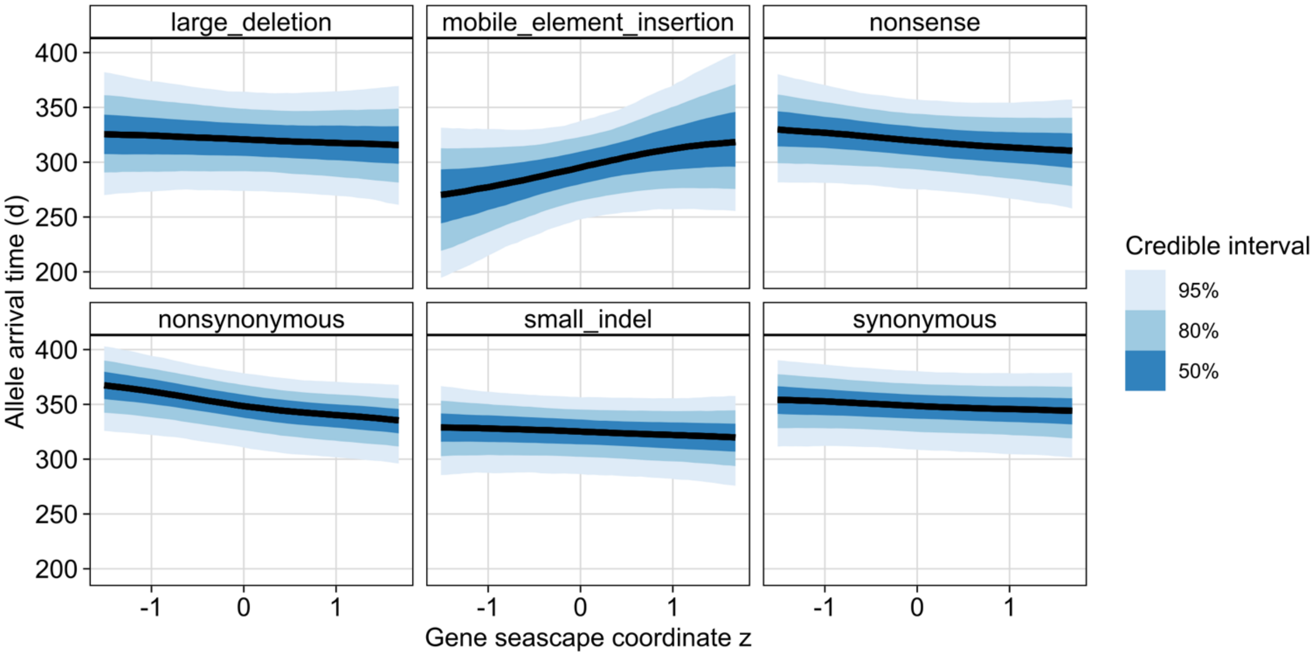


**Figure S10. Seascape coordinate predicts allele arrival time across certain mutation classes.** Posterior predictions from a cumulative-logit ordinal mixed model predicting allele arrival time (binned by sampling interval) from seascape coordinate *z*, shown separately for each mutation class. Lines show posterior medians and ribbons indicate 50%, 80%, and 95% credible intervals. Models include sample-level random intercepts.

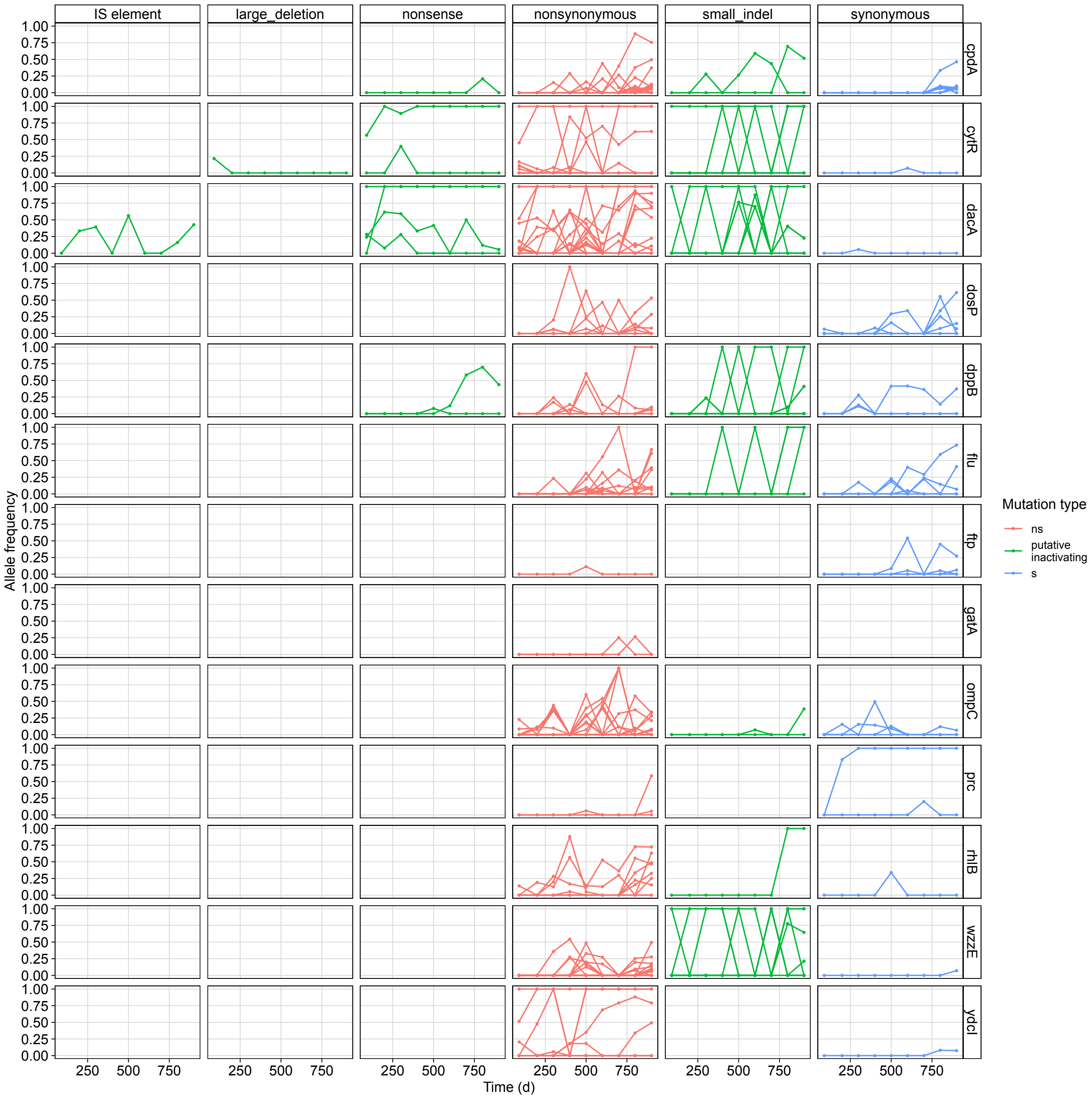


**Figure S11. Locus-specific mutation frequency trajectories across repeatedly mutated genes in feast/famine evolution.** Allele frequency time series are shown for recurrently mutated loci across replicate evolving populations. Row correspond to genes and columns to mutation classes. Each trajectory represents a unique allele, with alleles from replicate populations combined in each panel. Colors group alleles into broad effect classes (nonsynonymous, synonymous, and putatively inactivating) as indicated in the legend.
